# The Kinesin-5 Tail Domain Directly Modulates the Mechanochemical Cycle of the Motor for Anti-Parallel Microtubule Sliding

**DOI:** 10.1101/713909

**Authors:** Tatyana Bodrug, Elizabeth Wilson-Kubalek, Stanley Nithianantham, Alex F. Thompson, April Alfieri, Ignas Gaska, Jennifer Major, Garret Debs, Sayaka Inagaki, Pedro Gutierrez, Larissa Gheber, Richard McKenney, Charles Sindelar, Ronald Milligan, Jason Stumpff, Steven Rosenfeld, Scott T. Forth, Jawdat Al-Bassam

## Abstract

Kinesin-5 motors organize mitotic spindles by sliding apart anti-parallel microtubules. They are homotetramers composed of two antiparallel dimers placing orthogonal motor and tail domains at opposite ends of a bipolar minifilament. Here, we describe a regulatory mechanism, involving direct binding of the tail to motor domain and reveal its fundamental role in microtubule sliding motility. Biochemical analyses reveal that the tail down-regulates microtubule-activated ATP hydrolysis by specifically engaging the motor in the nucleotide-free or ADP-bound states. Cryo-EM structures reveal that the tail stabilizes a unique conformation of the motor N-terminal subdomain opening its active site. Full-length kinesin-5 motors undergo slow motility and cluster together along microtubules, while tail-deleted motors exhibit rapid motility without clustering along microtubules. The tail is critical for motors to zipper together two microtubules by generating substantial forces within sliding zones. The tail domain is essential for kinesin-5 mitotic spindle localization *in vivo*, which becomes severely reduced when the tail is deleted. Our studies suggest a revised microtubule-sliding model, in which tail domains directly engage motor domains at both ends of kinesin-5 homotetramers enhancing stability of the dual microtubule-bound states leading to slow motility yet high force production.

## Introduction

Microtubules (MTs) form tracks for the active transport of vesicles and macromolecules inside eukaryotic cells, generate pulling forces during assembly of mitotic spindles, and promoting the alignment and segregation of chromosomes (Goshima and Scholey, 2010; Vale, 2003). Fourteen kinesin motor subfamilies utilize MTs as tracks for these diverse functions (Vale, 2003). Among them, kinesin-5 motors represent a unique and highly conserved subfamily that is essential for mitotic spindle assembly during metaphase, and for spindle elongation during anaphase (Kashina et al., 1996). In contrast to the majority of dimeric kinesin classes, kinesin-5 motors adopt a conserved bipolar homotetrameric organization, composed of two dimeric subunits folded in an antiparallel arrangement mediated by the assembly of a 60-nm long central minifilament (Acar et al., 2013; Kashina et al., 1996; Scholey et al., 2014; Singh et al., 2018). Through this conserved bipolar organization, kinesin-5 motors promote MT crosslinking and their sliding apart during mitotic spindle assembly and elongation. This activity can recapitulated *in vitro* with purified kinesin-5 orthologs from a variety of species (Kapitein et al., 2008; Kapitein et al., 2005; van den Wildenberg et al., 2008).

Metazoan kinesin-5 orthologs such as *D. melanogaster* KLP61F or human Eg5 undergo slow plus-end directed motility especially during antiparallel MT sliding (Kapitein et al., 2008; Kapitein et al., 2005; Shimamoto et al., 2015; van den Wildenberg et al., 2008). In contrast, yeast kinesin-5 orthologs, such as Cin8, Kip1 and Cut7 uniquely undergo minus-end directed motility as single motors and reverse direction toward MT plus-ends upon clustering into multi-motor assemblies along single MTs, or during antiparallel MT sliding (Edamatsu, 2014; Fridman et al., 2013; Gerson-Gurwitz et al., 2011; Roostalu et al., 2011; Shapira et al., 2017). The conserved plus-end directed MT sliding activity is essential for mitotic spindle assembly by generating forces exerted on MTs emanating from opposite spindle poles during metaphase and stabilizing the characteristic bipolar spindle organization (Brust-Mascher et al., 2009; Forth and Kapoor, 2017; Subramanian and Kapoor, 2012; Wang et al., 2014). The MT sliding activity is critical for the elongation of mitotic spindles at the midzone region during anaphase (Goshima and Scholey, 2010). Defects in mammalian kinesin-5 orthologs or their inactivation via inhibitory compounds, such as monastrol, result in monopolar spindles by disruption in the balance of mechanical forces within the spindle (Goshima and Scholey, 2010; Goshima et al., 2005). These inhibitory compounds aided in elucidating the fundamental functions of kinesin-5 in mitosis and were suggested to be of therapeutic value in treating rapidly dividing cancer cells (Kwok et al., 2006; Mayer et al., 1999; Owens, 2013).

Each Kinesin-5 motor consists of a conserved organization including: an N-terminal motor domain, α-helical neck and bipolar assembly regions, and a C-terminal tail domain. The motor domain is connected via a neck-linker to a dimerizing neck α- helical coiled-coil (Turner et al., 2001; Valentine et al., 2006a; Valentine et al., 2006b). The parallel α-helical coiled-coil neck forms a part of the 60-nm central antiparallel homotetrameric α-helical minifilament (Acar et al., 2013; Kashina et al., 1996). At the center of this minifilament is a 27-nm antiparallel four α-helical bundle termed the bipolar assembly (BASS) region (Scholey et al., 2014). The kinesin-5 BASS tetramer orients two parallel neck coiled-coils and their associated motor domains to be off-set through a 100°-lateral rotation with respect to each other, potentially mediating their preference in binding and sliding two antiparallel MTs (Scholey et al., 2014). An extended section of unknown structure connects the C-terminal end of the BASS to the tail domain, which resides in close proximity to the motor domains of the antiparallel subunits (Acar et al., 2013). Thus, each kinesin-5 end consists of twin tail and twin motor domains originating from two sets of antiparallel folded dimeric subunits that emerge in close proximity at each end of the bipolar homotetramer (Acar et al., 2013; Scholey et al., 2014).

The kinesin-5 tail domain contains a conserved BimC box, which is a consensus motif that is phosphorylated by mitotic cyclin-dependent kinases (Blangy et al., 1995; Sharp et al., 1999). Mitotic phosphorylation at the BimC box induces kinesin-5 motors to concentrate along the mitotic midzone, where they promote the elongation of the mitotic spindle during late anaphase by sliding apart antiparallel MTs (Sharp et al., 1999). However, a role for this phosphorylation in the regulation of kinesin-5 activity remains unknown. Studies of the *S. cerevisiae* yeast ortholog, Cin8, show the tail domain is essential for kinesin-5 function, where its deletion leads to a lethal mitotic arrest phenotype in the absence of Kip1(Hildebrandt et al., 2006). The tail domain of the *Xenopus* Eg5 was suggested to form a secondary MT binding site during MT sliding motility, yet the function of the kinesin-5 tail-MT interaction remains unclear (Weinger et al., 2011). Despite extensive structural, kinetic and functional analyses of kinesin-5 orthologs, the origin of their highly conserved MT sliding activity and its relation the conserved tetrameric organization remains poorly understood.

Here, we describe a mechanism for the tail to motor domain regulation within homotetrameric kinesin-5 motors and its fundamental role during MT sliding motility. Using biochemical methods, we show the kinesin-5 tail domain down-regulates MT-activated ATP hydrolysis through binding and stabilizing the MT-bound motor domains in the ADP or nucleotide-free states. Cryo-EM structures reveal that the tail stabilizes the open conformation of the motor by binding its N-terminal subdomain via the α0-helix element at its tip. We show that human Eg5 motors undergo very slow motility and form clusters along single MTs, whereas Eg5 motors with their tails deleted undergo rapid motility without clustering along single MTs. Single motor spiking MT sliding assays show that Eg5 motors undergo slowed unidirectional motility within active MT sliding zones, whereas tail-deleted Eg5 motors undergo rapid motility with frequent switches in direction along either antiparallel MT within sliding zones. Optical trapping and MT sliding assays reveal that the tail is essential for Eg5 motors to zipper two MTs into sliding zones by producing substantial pushing forces. In contrast, tail deleted Eg5 motors exhibit severe defects in zippering two MTs leading to a poor capacity to generate forces. Tail deletion leads to a loss of Eg5 mitotic spindle localization in mammalian cells, while retaining MT binding activity. Our studies suggest a revised kinesin-5 MT sliding model in which the tail domain down-regulates MT-activated ATP hydrolysis at each end of the homotetramer, enhancing force production for sliding apart MTs, by stabilizing the dual MT-bound states along both sliding MTs.

## Results

### The kinesin-5 tail domain downregulates MT activated ATP hydrolysis by binding the motor domain in the ADP or nucleotide-free states

To better understand the role of the tail domain on kinesin-5 function, we first evaluated the biochemical effect of the kinesin-5 tail domain on the MT stimulated ATP hydrolysis by the kinesin-5 motor domain. To circumvent the added complexity of MT network formation due to the MT sliding activity of homotetrameric kinesin-5 motors (Figure 1A), we studied MT-activated ATP hydrolysis for constructs of motor-neck-linker domains (termed “motor”; residues 1-356) in the presence and absence of the isolated tail (termed “tail”; residues 913-1056) or a fusion construct in which the motor-neck-linker is linked at its C-terminus to the tail (termed “motor-tail fusion”) via an 8-residue linker (Figure 1B-D). We studied both human-Eg5 and Dm-KLP61F constructs to determine the conservation across these two well-studied orthologs (Table 1; Figure 1B-D; Figure S1C-D). The MT-activated ATP hydrolysis rate was measured by titrating an increasing concentration of MTs to the motor, a mixture of motor and tail, or motor-tail fusion (Materials and Methods; Figure 1B-D). The KLP61F motors robustly hydrolyzed ATP in response to increasing amounts of MTs (k_cat_ 7.1 s^−1^ and K_m_ 680 nM; Figure 1B; Table 1). The addition of equimolar KLP61F tail to a mixture of the motor and MTs leads to two-fold decrease in k_cat_ (3.5 s^−1^ vs 7.1 s^−1^) but produces little change in K_m_ (756 nM vs 680 nM) suggesting that the tail does not competitively interfere with motor-MT binding but rather modulates MT-activated ATP hydrolysis (Figure 1C). The motor-tail fusion exhibited a 2.2-fold decrease in k_cat_ compared to the motor alone (3.4 s^−1^ vs 7.1 s^−1^) and a 15-fold decrease in K_m_ compared to that for the motor alone (39 nM vs 680 nM) (Figure 1D). We observed similar behavior for human Eg5 constructs. The MT activated ATP hydrolysis measurements reveal that the Eg5 tail down-regulates the MT activated motor ATP hydrolysis rate by 32% compared to that for Eg5 motor alone (7.6 s^−1^ vs 5.3 s^−1^)(Table 1; Figure S1G). These data reveal that the kinesin-5 tail domain down regulates the MT activated ATP hydrolysis of the motor domain in either the isolated or fused configurations.

**Figure 1:**
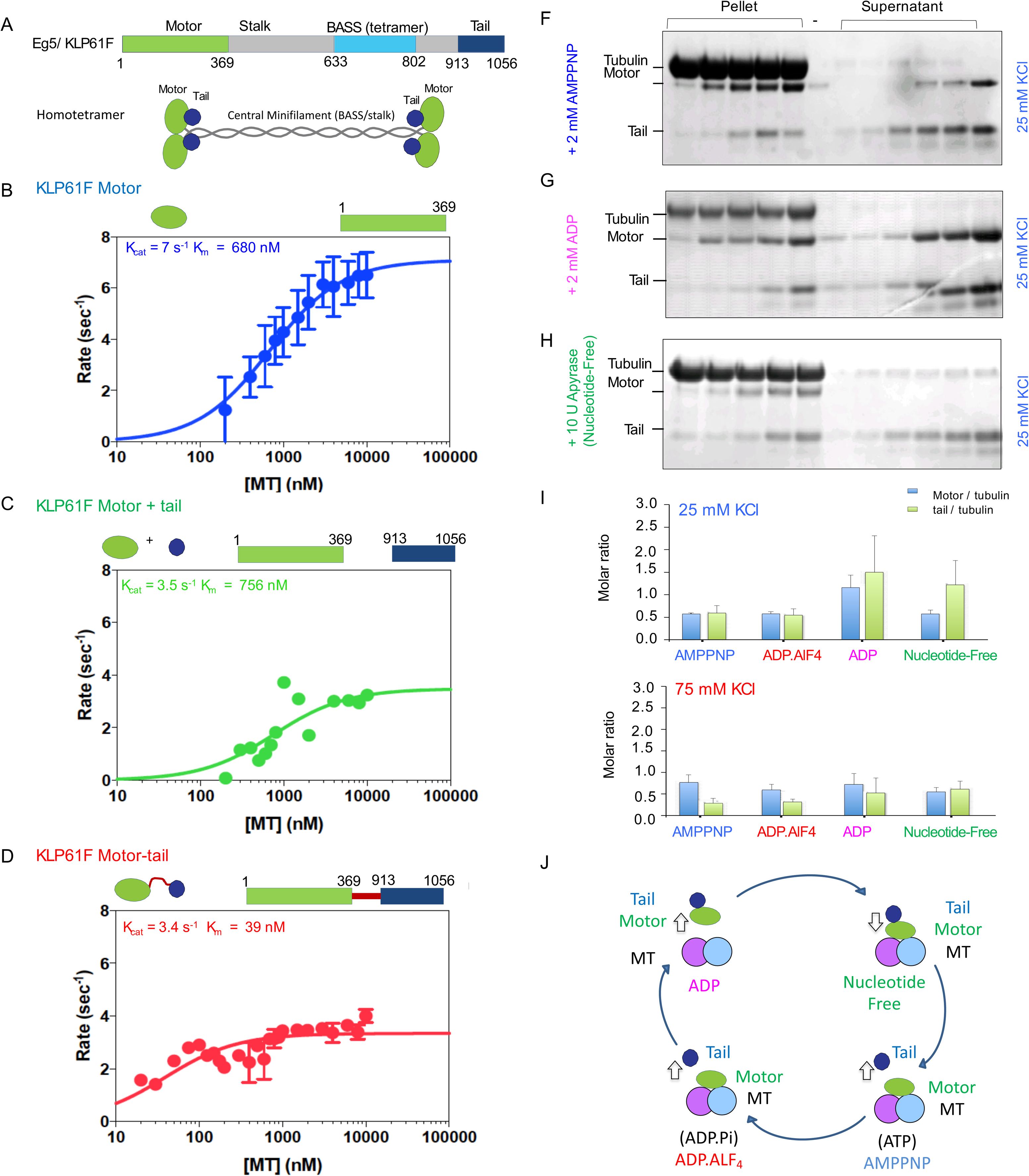
The kinesin-5 tail domain inhibits the motor domain MT-activated ATPase through stabilizing the MT bound nucleotide-free state. A) Top, the domain organization of kinesin-5 orthologs, Dm KLP61F and Hs FL-Eg5, revealing their conserved N-terminal domain, central BASS domain and C-terminal tail domain. Bottom, domain homotetrameric organization for kinesin-5 motors. B) Steady state ATP hydrolysis for KLP61F motor with increasing MT concentration. C) Steady state ATP hydrolysis for equimolar KLP61F motor + tail constructs with increasing MT concentration. D) Steady state ATP hydrolysis for KLP61F motor-tail fusion with increasing MT concentration. E) MT co-sedimentation assays of the KLP61F motor (motor) and tail domain (tail) with MTs (MT) in the presence of non-hydrolysable analog, 2 mM AMPPNP at 25 mM KCl. Co-sedimentation at 25 mM KCl compared to 75 mM KCl are shown in Figure S2B. F) MT co-sedimentation assays of the KLP61F motor (motor) and tail domain (tail) with MTs (MT) in the presence of the 2 mM ADP at 25 mM KCl. Co-sedimentation at 25 mM KCl compared to 75 mM KCl are shown in Figure S2C. G) MT co-sedimentation assays of the KLP61F motor (motor) and tail domain (tail) with MTs (MT) in the nucleotide-free state, achieved by adding 10 U of Apyrase. Co-sedimentation at 25 mM KCl compared to 75 mM KCl are shown in Figure S2D. H) Quantitative densitometry of data in Figure S1A-D reveals the molar ratios of motor to tubulin monomer (blue) and tail to tubulin dimer (green). The motor-to tubulin ratio observed at 75 mM KCl is roughly 0.5 and the amount of the tail bound to the motor increases in the ADP and Nucleotide-free states increases in the 25 mM KCl compared to the 75 mM KC. The amounts of tail bound to motor remain low in the AMPPNP and ADP.AlF4 states compared to higher amount in the ADP and nucleotide-free states.

To understand the biochemical basis for the kinesin-5 tail mediated down-regulation of the motor MT-activated ATP hydrolysis, we reconstituted binding of the motor and tail constructs to Paclitaxel-stabilized MTs in four nucleotide state conditions and analyzed the binding using MT co-sedimentation assays (Figure 1F-I; Figure S1A-F). These conditions trap the motor in distinct nucleotide states, mimicking each step of the kinesin ATP hydrolysis cycle (Figure 1F-I; Figure S1A-F). We compared the binding activities at two ionic strength conditions (25 and 75 mM KCl), for which our motility assays indicate two different modes of tail dependent motility regulation (see below; Figures 3–4). The KLP61F tail and motor each bound MTs in isolation in the presence of 2 mM ADP, as has been shown previously (Figure S1E-F)(Weinger et al., 2011). The molar ratio of the bound tail to MTs is around ~1 tail per αβ-tubulin dimer site at 25 mM KCl and it decreased to ~0.5 at 75 mM KCl (Figure S1F). MT co-sedimentation of motor and tail in the presence of 2 mM AMPPNP or ADP.AlF4 revealed that the tail dissociated from MTs, while the motor bound MTs to saturation (Figure 1F; Figure S1A-B). In these conditions, the high affinity of the motor for MTs displaced the tail, which remained mostly in the supernatant. Even at the lowest molar ratio of motor to MTs, where there is an abundance of unoccupied MT lattice sites, we observe that the tail does not bind MTs and does not compete with the motor for MT binding sites. MT co-sedimentation in the presence of 2 mM ADP, reveal the motor binds MTs with low affinity in a concentration and ionic strength dependent manner, where its affinity is increased at 25 mM KCl compared to 75 mM KCl (Figure 1G; Figure S1C). At 25 mM KCl and 2 mM ADP, a higher amount of motor is bound to MTs, leading to higher amounts of tail being recruited into the pellet, approaching a molar ratio of ~1 motor per αβ-tubulin lattice site (Figure 1G, I). At 75 mM KCl, we observe a concomitant decrease in both the motor and tail to molar ratios of ~1.0 to ~0.5 of MT lattice sites (Figure 1I). Thus, MT co-sedimentation of the tail depends on the amount of MT-bound motor recruited to the pellet, which differ at 25 and 75 mM KCl, rather than direct binding to MTs (Figure 1G; Figure S1C). MT co-sedimentation in the presence of apyrase (1U/ml) reveals a high affinity of the motor in the nucleotide free state in both 25 and 75 mM KCl (Figure 1H). In this condition, a higher amount of tail is recruited and it correlated with the amount of the motor bound MT fraction (Figure 1I). This suggests that the tail affinity for the MT bound motor in the nucleotide-free or ADP states is high (Figure 1I). Quantitative densitometry analyses indicate the molar ratios of tail recruited to the MT bound fraction are the highest in the ADP and nucleotide-free state in contrast to the AMPPNP and ADP.AlF4 states (Figure 1I). We also studied analogous human Eg5 motor and tail constructs in above three nucleotide states revealing essentially similar patterns of findings (Figure S1H-J). Thus, together these data reveal that the ATP and ADP.Pi states of the kinesin-5 motor domain exhibits a high binding affinity for MTs, but a low affinity for the tail domain, and displace the tail from MTs. Whereas in ADP and nucleotide-free states of the motor exhibit increased affinity for the tail, recruiting the tail to the MT bound fraction despite the difference in the motor MT affinity in these two states.

### Cryo-EM structure of the motor-tail interface reveals the tail domain stabilizes an open conformation of the motor domain active site

We used cryo-electron microscopy (cryo-EM) to investigate the kinesin-5 tail-motor interface and its role in down-regulating ATP hydrolysis. New and previously described image analysis strategies were used to calculate and refine structures of MTs decorated with the KLP61F motor in the AMPPNP state and the KLP61F motor-tail in the nucleotide-free state to ~4.4 Å (Figure 2A-F; Figure S2A-C; Table 2; see materials and methods). The AMPPNP kinesin-5 motor structure revealed the conserved kinesin fold with two longer and reorganized class specific L6 and L8, which forms part of the MT-interface (Figure S2E-F). The map is very similar to the recently-determined *S. pombe* and *U. maydis* kinesin-5 motor domain structures (von Loeffelholz and Ann Moores, 2019; von Loeffelholz et al., 2019).

**Figure 2:**
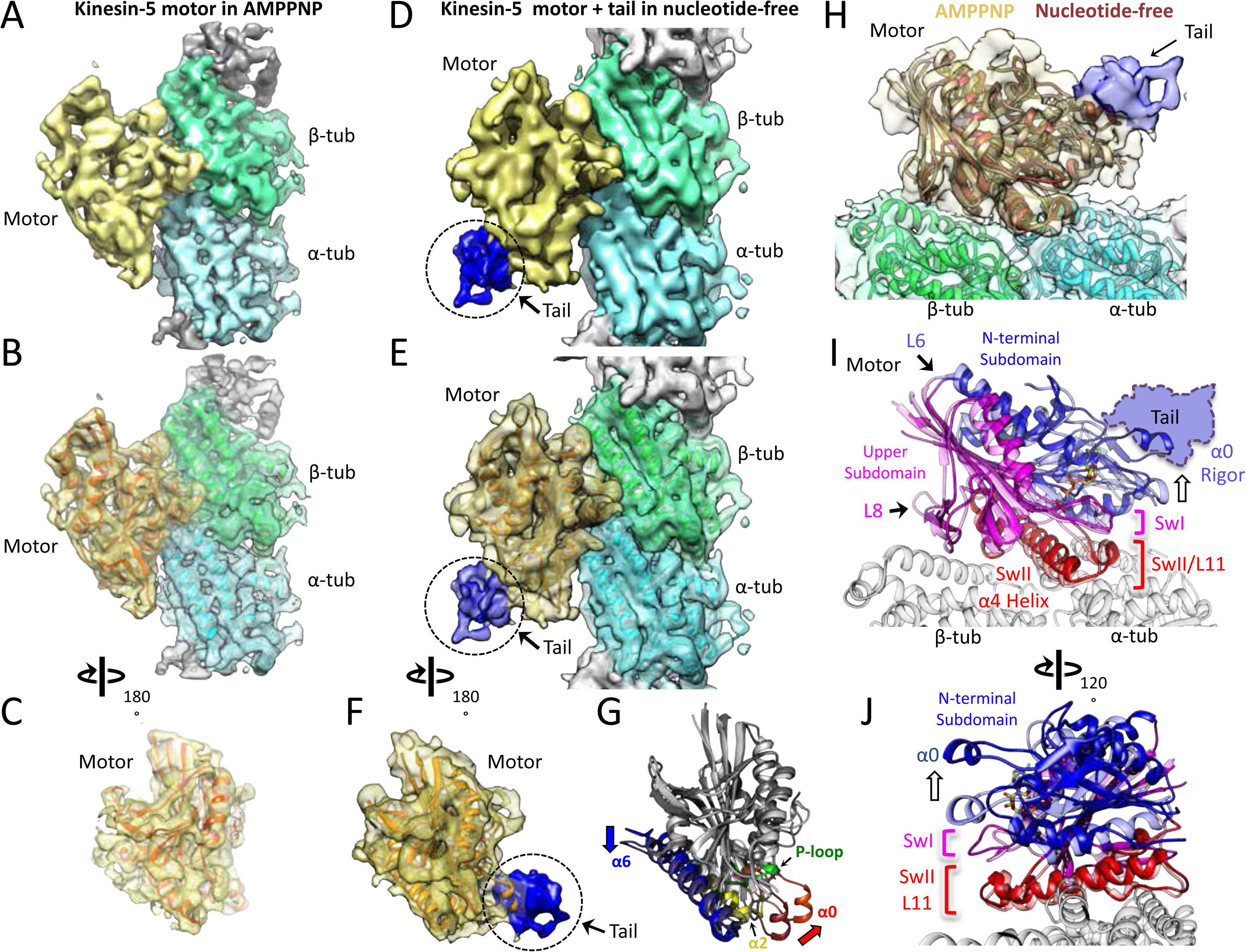
The kinesin-5 tail domain engages the motor domain through direct interface near α0 helical hairpin in the nucleotide-free state. A-C) A: Side view of 4.4-Å cryo-EM structure of KLP61F motor domain decorated MT unit in the AMPPNP state. A single kinesin-motor bound αβ-tubulin unit is shown. The segmented motor domain (yellow) and α-tubulin (cyan) and β-tubulin (green) densities are sown. B: de novo built KLP61F motor domain model in the AMPPNP state (red) displayed within the motor domain density. The αβ-tubulin dimer model fitted into the αβ-tubulin density (green and cyan). C: Top end view of the kinesin-5 motor density map with the MT density computationally removed. D-F) D: Side view of 4.0-Å cryo-EM structure of KLP61F motor and tail domains decorated MTs in the nucleotide-free state obtained using well-established and refined with new strategies (Figure S2A-C; see materials and methods). The kinesin-motor + tail bound a single αβ-tubulin unit is shown. Segmentation of motor domain (yellow), tail domain is shown (blue), α-tubulin (cyan) and β-tubulin (green). E: de novo built KLP61F motor domain model in the nucleotide-free state (orange) displayed within the motor domain density and the tail density (dark blue). The αβ-tubulin dimer fitted into the αβ-tubulin density (green and cyan). F: Top end view of the docked kinesin-5 motor and tail density with the MT density computationally removed. G) Conformational transition of KLP61F motor domain from nucleotide-free (light grey) to AMPPNP (dark grey). The elements that undergo the most change in colors: α6 (blue), α2 (yellow), p-loop (green) and α0 helix (red). I-J) Two views of the KLP61F AMPPNP and nucleotide-free motor domain models describing the movements of with N-terminal subdomain (deep blue, nucleotide-free; light blue, AMPPNP) and Upper subdomain (deep pink, nucleotide-free; light pink, AMPPNP) around the α4-helix, L11 switch II (swII) MT bound subdomain (deep red, nucleotide-free; light red, AMPPNP). The switch I (swI) changes conformation in response to ATP binding. The N-terminal subdomain rotation leads to 20° rotation of α0-helix closer to the MT surface in the AMPPNP state. The tail binds α0-helix in the nucleotide-free state. H) Side view of the KLP61F motor domain maps in AMPPNP (red) compared the nucleotide-free state model (orange) docked into the nucleotide free-state motor density with the tail density shown in blue

Class averages for the motor MT-decorated segments in the two conditions clearly reveal repeating motor-densities decorating the MT lattice sites (Figure S2D). Additional density is observed on the motor domain in the nucleotide-free motor-tail decorated MTs. Despite the size of the tail (80-100 residues), the density attributable to it in the 3D maps is small and accounts for only about one third this mass (Figure S2B-C, G). This suggests that a large proportion of the tail is either unstructured, or forms a flexibly attached, separate domain; a major part of the tail region is rendered invisible after averaging. Furthermore, the density seen cannot be interpreted in terms of secondary structure as there is a substantial drop in the resolution to ~8-Å at the motor-tail interface (Figure S2B-C; Video S1). In keeping with the biochemical data, we do not observe any direct, ordered interaction between the tail density and the MT lattice. Overall, the data strongly suggest that the tail is interacting loosely with the underlying motor domain, binding to a small region on the motor, located on its MT minus end facing side (Figure 2D-F). Comparison of the KLP61F motor AMPPNP structure with the motor-tail nucleotide-free structure reveals conformational change within the motor domain (Figure 2D-F), and suggests how elements of the nucleotide-free state form a binding site for the tail (Figure 2C). The tail binds an exposed hairpin loop at the end of the α0 helix in the motor nucleotide-free state (Figure 2E-F). The other end of the α0 helix is connected to the phosphate binding loop (p-loop) (Figure 2G), thus allowing possible feedback between the nucleotide pocket and the tail binding site.

Model building for both the ATP-like and nucleotide-free motor domain structures reveals conformational changes that regulate the affinity of the tail within the motor domain (Figure 2C,F; Video S1). Structurally, the kinesin motor domain can be divided to three subdomains: the N-terminal subdomain, the upper subdomain, and the MT-binding subdomain (Figure 2FH-I). The MT-bound subdomain consists of the α4-helix L11 and L12, as described for kinesin-1 (Figure 2I-J; Figure S2E-F; Video S1)(Cao et al., 2014; Shang et al., 2014). As in kinesin-1, the N-terminal subdomain rotates around the MT subdomain in the kinesin-5 nucleotide-free state, leading to a reorganization of the MT minus-end facing end of the motor (Figure 2H-I; Figure S2E-F). In the nucleotide-free state, N-terminal subdomain (blue) rotates upward with respect to the upper subdomain (pink) and the MT bound α4 helix (switch-II) and L11 (red), opening the switch I loop in active site (Figure 2H-J). Compared to the AMPPNP state, the N-terminal subdomain in the nucleotide-free state rotates upward by 20° leading the α0-helix, which lies at its extreme tip, to move upward by 10-Å from the MT surface (Figure 2H-J; Video S1). This N-terminal subdomain rotation repositions the switch I, switch II, and P-loops, leading to an open active site conformation (Figure 2H-J). While in the AMPPNP state, the α0-helix is positioned downward, 10-Å closer to the MT lattice, (Figure 2H-J) due to ATP binding and active site closure. Our structures suggest that tail domain binding to the N-terminal subdomain likely stabilizes the upward α0-helix conformation, inhibiting transition of the P-loop, switch I//II from engaging ATP (Figure 2J; Video S1). Our models suggest that the tail stabilizes this upward N-terminal subdomain motor conformation, which likely down-regulates the ability of the active site to bind incoming ATP. This suggests that the motor-α0-helix may cycle into “on” and “off” states while binding the tail resulting in a slow ATP hydrolysis cycle.

### The tail domain down-regulates kinesin-5 motility velocity along single MTs

To examine the role of the tail to motor interface in kinesin-5 motility, we reconstituted the motility properties of homotetrameric human Eg5 motors along individual MTs. We studied two human Eg5 motor constructs that either contain or exclude the tail domain (Figure 3A). We purified full-length Eg5 (termed FL-Eg5, residues 1-1056), full-length Eg5 with a C-terminal GFP (termed FL-Eg5-GFP), and mutant Eg5 with the tail domain deleted with a C-terminal GFP (termed Eg5-Δtail-GFP, residues 1-912) (Figure 3A-B). These motors were expressed in insect cells and purified using StrepII affinity, followed by size exclusion chromatography (Figure S3A-B; see materials and methods). Deleting the tail domain does not alter the shape or the homotetrameric oligomerization of Eg5, as was described previously (Acar et al., 2013; Weinger et al., 2011)(Figure 3B; Figure S3A-B). We reconstituted FL-Eg5-GFP and Eg5-Δtail-GFP motility along GMPCPP or Paclitaxel-stabilized AlexaF-633 labeled MTs using TIRF microscopy assays (Figure 3A, lower panel). Both FL-Eg5-GFP and Eg5-Δtail-GFP exhibited robust and processive motility along MTs at 25 mM HEPES pH 7.5 containing 25 to 100 mM KCl conditions (Figure 3D-E; Video S2). This processive motility is highly homogenous in contrast to the non-processive diffusive motility that has been seen at 80 mM PIPES pH 6.8 (BRB-80) with a range of 0-100 mM KCl conditions (Kapitein et al., 2008; Kapitein et al., 2005)(data not shown). These observations suggest that the buffer composition, which influences pH and ionic strength, is very critical for observing robust Eg5 motor motility along single MTs.

**Figure 3:**
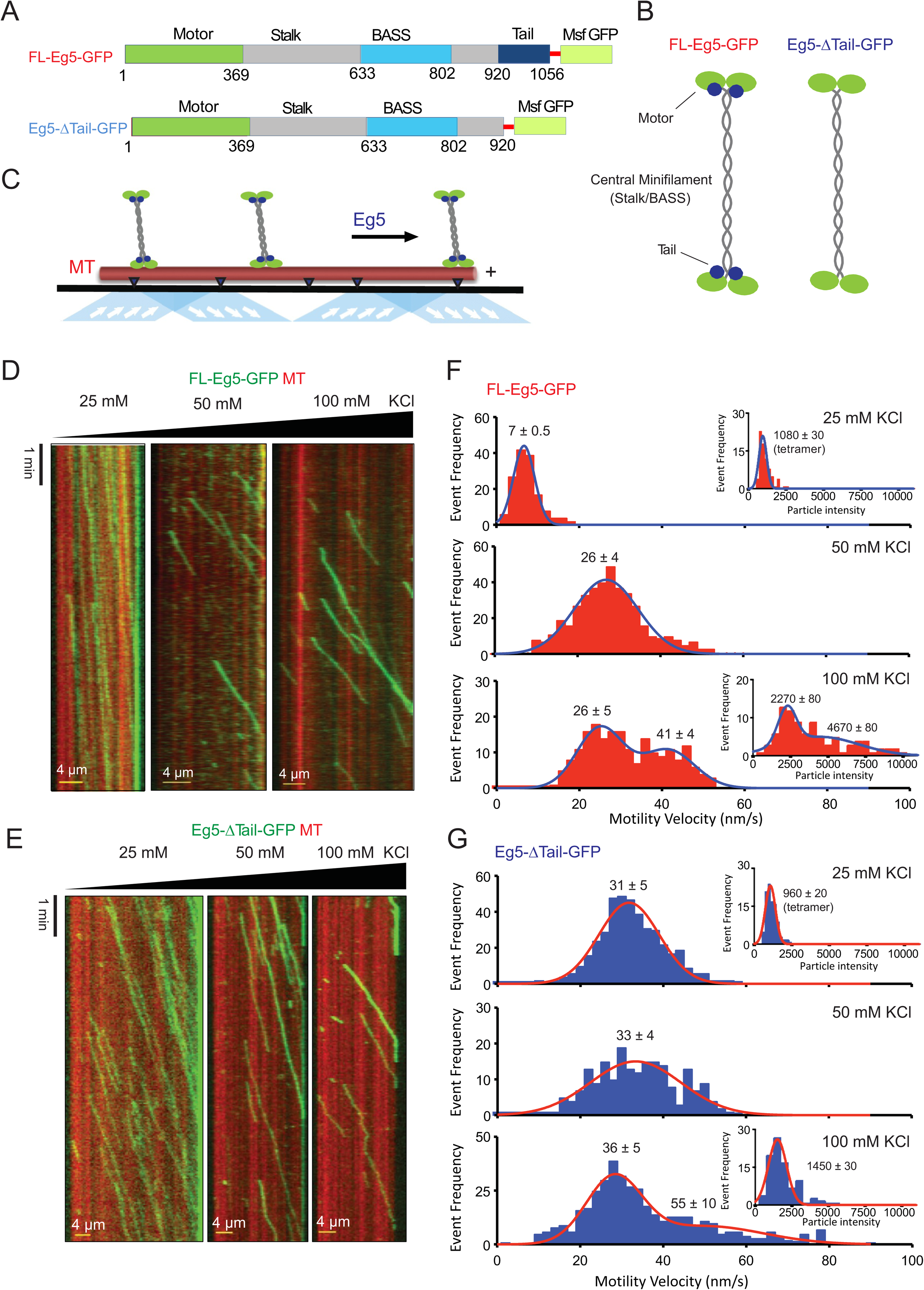
The tail domain down-regulates the motility velocity of homotetrameric kinesin-5 motors along MTs. A) Constructs of human Eg5 used in reconstitution studies. Top panel, domain organization of FL-Eg5-GFP. Second panel, Domain organization of Eg5-Δtail-GFP with its tail domain deleted (residues 913-1058) with the C-termini fused to monomeric superfolder (msf)-GFP. B) The homotetrameric Eg5 organizations for the two constructs FL-Eg5-GFP and Eg5-Δtail-GFP, as described previously (Acar et al 2013). C) scheme for TIRF microscopy of Eg5 motors (green) undergoing motility along single surface anchored AlexaF-633 and biotin labeled MTs via neutravidin-biotin attachment (red). D) Kymographs of FL-Eg5-GFP motor motility along MTs in 25, 50 and 100 mM KCl pH 7.5 condition. kymographs in dual color showing MTs (red) and GFP channels (green) are shown. Left panel, FL-Eg5-GFP motors undergo extremely slow motility at 25 mM KCl motor and their particle intensities are uniform. Middle panel, FL-Eg5 GFP undergoes motility with increased velocity and exhibits variation in intensity. Right panel, FL-Eg5-GFP motor undergo motility with higher velocity and exhibit bright and dim intensity particles at 100 mM KCl. Note FL-Eg5-GFP accumulate at MT plus-ends in 25 mM KCl and its plus-end residence decreases at 50 and 100 mM KCl. E) Kymographs of Eg5-Δtail-GFP motility along anchored MTs at 25, 50 and 100 mM KCl pH 7.5 condition. kymographs in dual color showing MTs (red) and GFP channels (green) are shown. Eg5-Δtail-GFP motors undergo motility at similar motility velocities in all conditions. Note the homogeneity in motor intensities for Eg5-Δtail-GFP and its rapid motility velocity at 25 mM KCl in contrast to the very slow motility of FL-Eg5-GFP. Motor intensities are uniform for both Eg5-Δtail-GFP at 25 mM KCl and remain mostly homogenously dim at 100 mM KCl. Note all Eg5-Δtail-GFP accumulate at MT plus-ends in a salt dependent manner. F) Top panel, histogram for FL-Eg5-GFP motor particle velocity frequency distribution revealing that homogenous and very slow velocities 25 mM KCl. Middle panel, histogram for FL-Eg5-GFP motor velocity frequency distribution at 50 mM KCl. Bottom panel, bimodal velocity frequency distribution for FL-Eg5-GFP at 100 mM KCl. Two inset motile Eg5 fluorescence intensity distribution for motile particles shown for 25 and 100 mM kCl revealing the motors are homogenous homotetramers at 25 mM KCl, but assemble into clusters of dimers to tetramers of homotetramers at 100 mM KCl. G) Histogram for Eg5-Δtail-GFP motor particle velocity frequency distribution (blue) 25, 50 and 100 mM KCl condition. Top panel at 25 mM KCl, Eg5-Δtail-GFP motors undergo only rapid motility velocity Eg5-Δtail-GFP motor which are three folds faster than FL-Eg5-GFP. Middle and right panels, motility velocity remains rapid in 50 and 100 mM KCl. Inset panels show that Eg5-Δtail-GFP motor fluorescence intensity distribution indicates that motors remain mostly as single homotetramers at 25-100 mM KCl in contrast to Fl-Eg5-GFP.

We analyzed the motility properties of FL-Eg5-GFP and Eg5-Δtail-GFP motors using the 25 mM HEPES pH 7.5 conditions buffer, containing 25, 50 and 100 mM KCl. The solution ionic strength generally influences the motor landing rates along MTs (Kapitein et al., 2008). At 25 mM KCl, FL-Eg5-GFP exhibited active yet extremely slow motility velocities (7 nm/s; Table 3; Figure 3D, left panel; Video S3), which was roughly four times lower than the motility velocity of Eg5-Δtail-GFP (31 nm/s; Figure 3E, left panel; Video S2). Large ensembles of motility events revealed mono-disperse distributions for velocities of the two types of motor that fit single Gaussians with single average values (Figure 3F-G, top panels). At 50 mM KCl, FL-Eg5-GFP exhibited more than three-fold increase in velocity (24 nm/s) compared to motility at 25 mM KCl (Figure 3D, middle panel). In contrast, Eg5-Δtail-GFP exhibited a nearly identical velocity (35 nm/s; at 50 mM KCl, which is indistinguishable from what was observed at 25 mM KCl (Table 3; Figure 3G, middle panel). At 100 mM KCl, FL-Eg5-GFP motors exhibited two velocities (24 and 41 nm/s), with a bimodal distribution representing 60% and 40% of total, respectively (Figure 3D right panel; Video S3; Figure 3F, lower panel). While in 100 mM KCl, Eg5-Δtail-GFP motors exhibited two velocities (36 nm/s and 55 nm/s), which are 20% higher than FL-Eg5-GFP and with a similar bimodal distribution representing 85% and 15% of the total, respectively (Table 3; Figure 3D right panel; Video S3; Figure 3G, lower panel). In the 50 and 100 mM KCl conditions, fewer FL-Eg5-GFP and Eg5-Δtail-GFP motor landing events were observed compared to 25 mM KCl (Figure 3D-E, right panels). Thus, the tail domain significantly slows the motility velocity of Eg5 homotetramers at 25 mM KCl, a feature that was absent from Eg5-Δtail-GFP. Similarly, at 50 to 100 mM KCl, Eg5-Δtail-GFP exhibited consistently 20% higher velocities compared to FL-Eg5-GFP with bimodal distributions of slow and fast particles (Table 3).

We next explored the higher-order oligomerization for FL-Eg5-GFP and Eg5-Δtail-GFP motile particles and their propensity to form clusters based on the relative fluorescence intensities of motile GFP-fused motors (Figure 3F-G, inset graphs). Kymographs show that FL-Eg5-GFP motors in 25 mM KCl and Eg5-Δtail-GFP motors in 25 and 100 mM KCl exhibited a homogenous and low intensity distribution that was well fit by a single Gaussian (Figure 3D-E, left panels; Video S3; Table 3; Figure 3F,G insert graphs in top panels). In contrast, FL-Eg5-GFP particles exhibited both a bright and dim intensity distributions and a bimodal intensity distribution at 100 mM KCl (Figure 3D, right panel; Video S3) which can be fitted by two overlapping Gaussians, representing a high intensity that is four-fold higher than dim motile particles observed at 25 mM KCl, and lower intensity that is roughly two folds higher than the dim particles at 25 mM KCl (Figure 3F, insert graph in bottom panel; Table 3). Whereas at 100 mM KCl, Eg5-Δtail-GFP motile particles exhibited a narrow intensity distribution that closely matched Eg5-Δtail-GFP motile particle distributions at 25 mM KCl (Figure 3G, insert graph in bottom panel; Table 3). Thus, FL-Eg5-GFP and Eg5-Δtail-GFP undergo motility as homogenous particles, likely individual homotetramers, with little clustering at 25 mM KCl. At 100 mM KCl, FL-Eg5-GFP motors form clusters consisting of up to two to four homotetramer at 100 mM KCl, unlike the Eg5-Δtail-GFP, which remained as individual homotetramers under all conditions tested. In the higher ionic strength settings, the tail domain may mediate inter-homotetramer interactions leading to this clustering.

Analyses of FL-Eg5-GFP and Eg5-Δtail-GFP motility run lengths reveal that the tail influences the rate of processive motility by kinesin-5 homotetramers. At 25 mM KCl, both FL-Eg5-GFP and Eg5-Δtail-GFP motors are highly processive, exhibiting long runs that extend throughout the entirety of the imaging time, eventually concentrating at MT plus-ends and were thus were not quantified (Figure S3C; Table 3). In contrast at 50 mM KCl, we observe measurable run lengths for FL-Eg5-GFP and Eg5-Δtail-GFP motility events showing a similar range of 13-14 μm. At 100 mM KCl, FL-Eg5-GFP retained their highly processive long run lengths with average 13 μm, while Eg5-Δtail-GFP showed a 45% decrease in run lengths to an average of 8 μm (Figure 3C; Table 3). These data suggest that the kinesin-5 motility cycle becomes less processive in the absence of the tail in Eg5-Δtail-GFP motor, due to a decrease in motor MT lattice affinity.

### The Kiensin-5 tail domain is required for efficient assembly of anti-parallel MT overlap zones

To understand the impact of the tail-to-motor interaction on kinesin-5 MT sliding motility, we reconstituted Eg5 MT sliding *in vitro* using three-color TIRF assays in 25 mM HEPES, pH 7.5 with 25-50 mM KCl conditions (Figure 4A). In this assay, AlexaF-633 and biotin-labeled MTs were attached to PEG-treated glass surfaces via a neutravidin to biotin linkage (anchored MT, red), and then AlexaF-560 labeled MTs (free MT yellow) and FL-Eg5-GFP or Eg5-Δtail-GFP motors were added to reconstitute the crosslinking and sliding of free-MTs (yellow) along the anchored MTs (red) (Figure 4A). Using this assay, we observed that 10-20 nM FL-Eg5-GFP motors promote robust MT sliding. In these assays, FL-Eg5-GFP motors bind along the anchored MTs, crosslink free-MTs (yellow) then zipper them to form anti-parallel MT sliding zones (Figure 4A-left; Figure S4; Video S4). FL-Eg5-GFP motors undergo slow motility similar to those observed along single MTs (Figure 3), and concentrate within newly formed two-MT bundles producing sliding zones (green) (Figure 4B-left). Next, we studied how increasing the solution ionic strength (25-50 mM KCl) influences the FL-Eg5-GFP motor velocity within the MT sliding zones and its effect on the sliding motility of the free-MTs (Figure 4C; Table 4). At 50 mM KCl, FL-Eg5-GFP motors displayed a two-fold increase in motility velocity similar to the effect along single MTs (Figure 3). The change in the velocity of individual motors along the anchored MTs matched a two-fold increase in the sliding rate of the free-MTs along the anchored MTs (Figure 4C; Table 4). These data directly correlate the kinesin-5 motor velocity and specifically tail mediated motor down-regulation of the motor motility velocity with the velocity of anti-parallel MT sliding (Table 4; Figure 4C).

**Figure 4:**
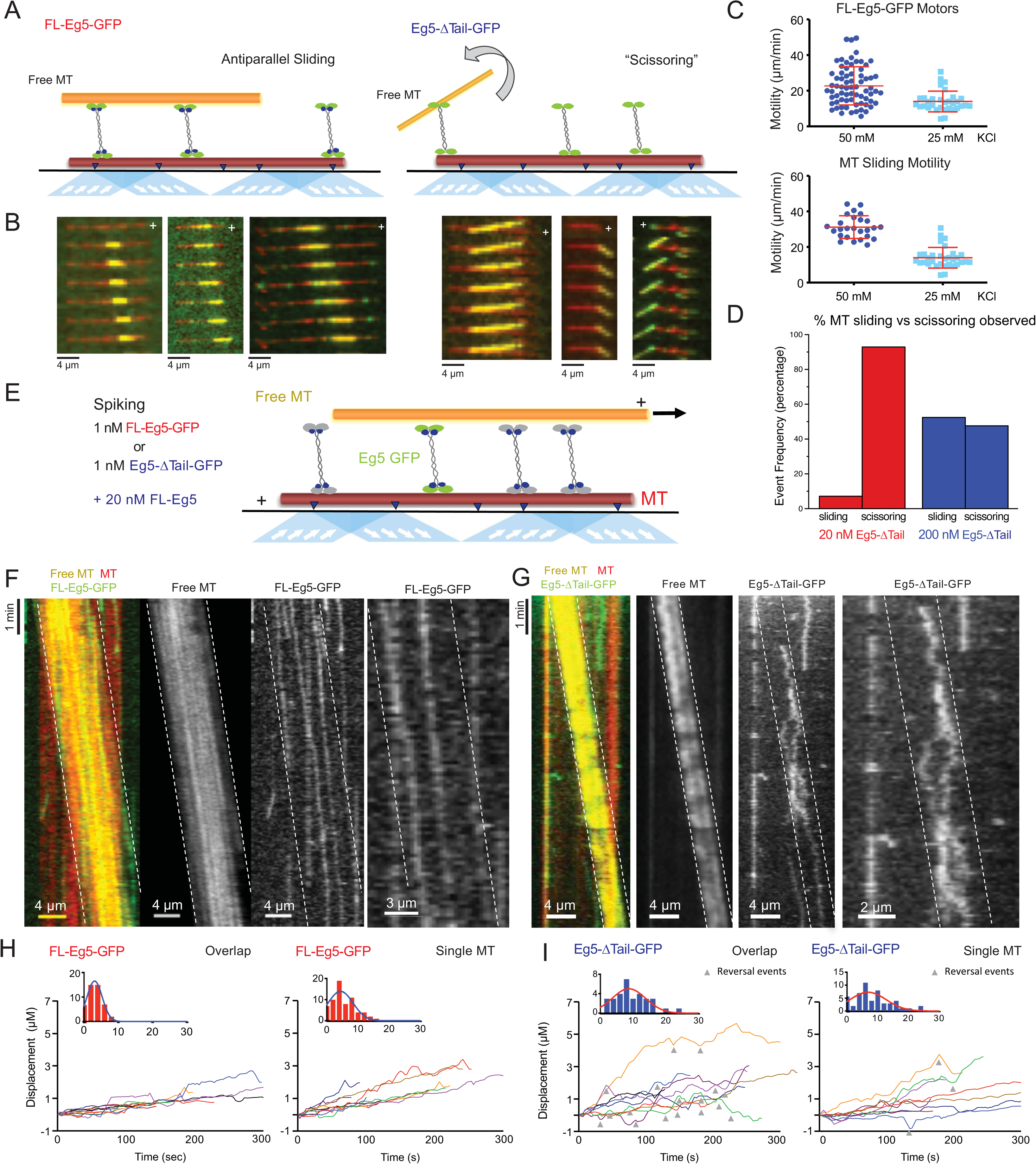
The kinesin-5 tail domain regulates the zippering of two sliding MTs via slow directional motility within the overlapping zones. A) Left, Scheme for TIRF microscopy for MT sliding assays where FL-Eg5-GFP motors (green) recruit free MTs (orange) along single surface anchored AlexaF-633 and biotin labeled MTs via neutravidin-biotin (red). Right, scheme for TIRF microscopy MT sliding where Eg5-Δtail-GFP mediates crosslinking of the free-MT (orange) without zippering them along the AlexaF-633 and biotin labeled MTs via neutravidin-biotin (red). Their activity leads to scissoring motility. B) Montages of different types of MT sliding events. left panels, FL-Eg5-GFP mediates MT sliding. Right panels, Eg5-Δtail-GFP mediates MT crosslinking but does not mediate zippering MTs leading to scissoring events. C) The influence of ionic strength on FL-Eg5-GFP MT sliding activity. Top panel, FL-Eg5-GFP motor particle motility velocities (top panel) and free MT sliding velocity (lower panel). D) The proportion of MT sliding and scissoring events in percentage of total observed for experiments at 20 and 200 nM Eg5-Δtail-GFP, respectively. The increase in Eg5-Δtail-GFP leads to higher proportion of MT sliding compared to scissoring events (N=70-100 sliding/scissoring events for each condition). E) Scheme for motor spiking during MT sliding assay with 20 nM FL-Eg5 (non-fluorescent, grey). 1 nM FL-Eg5-GFP or 1 nM Eg5-Δtail-GFP (green) were added to visualize single motors undergo motility along the anchored MTs (red) and transition into the MT sliding zone with both free (yellow) and anchored MT (red). F) Examples kymographs of MT sliding spiking assays with 1 nM FL-Eg5-GFP in the presence of 20 nM FL-Eg5 (unlabeled). Three color kymographs are shown including the overlay (right panel), the free MT sliding (middle panel, yellow) being slide apart in the presence of single FL-Eg5-GFP motors (middle panel, green). Extreme right panel, is magnified view revealing the slow and directional motility of FL-Eg5-GFP motors. G) Examples kymographs of MT sliding spiking assays with 1 nM Eg5-Δtail-GFP in the presence of 20 nM FL-Eg5 (unlabeled). Three color kymographs are shown including the overlay (right panel), the free MT sliding (middle panel, yellow) being slide apart in the presence of single Eg5-Δtail-GFP motors (middle panel, green). Extreme right panel, is magnified view. Note FL-Eg5-GFP motors undergo bi-directional motility with stochastic switching along either of the two anti-parallel MTs in the MT sliding zones, H) Single motor track quantitation of 1 nM FL-Eg5-GFP motor motility inside the MT sliding zone (left, overlap) and along the anchored MT (right, single). Note FL-Eg5-GFP undergoes slow motility in general but its velocity decreases even further in the MT sliding zone. I) Single motor track quantitation of 1 nM Eg5-Δtail-GFP motor motility inside the MT sliding zone (left, overlap) and along the anchored MT (right, single). Note Eg5-Δtail-GFP undergoes rapid motility in both zones, but its motility switches direction (reversals marked by grey arrow heads) particularly within the overlap region of the MT sliding zone.

We next studied the Eg5-Δtail-GFP in the MT sliding assays at similar concentrations (10-20 nM). Strikingly, the Eg5-Δtail-GFP motors only mediated crosslinking of the free-MT at focal points, and were unable to completely zipper the free MT along its length to the anchored MT (Figure 4A, right; Figure 4B right; Video S4). This poor MT zippering defect exhibited by the Eg5-Δtail-GFP motors leads the free MTs to “scissor” along the anchored MTs due to Brownian motion around the crosslinking point (Figure 4B-right; Video S4). The scissoring behavior occurs most frequently near or at MT plus-ends, where the Eg5-Δtail-GFP motors concentrate (Figure 4B, lower right, second and third panels). We next studied the effect of increasing the Eg5-Δtail-GFP motor on the formation of MT sliding zones. A ten-fold higher concentration of Eg5-Δtail-GFP (200 nM) lead to a higher proportion of MT sliding events compared to scissoring events (Figure 4D). Quantitative analyses revealed that at 20 nM Eg5-Δtail-GFP, only 6% of MT crosslinking events transitioned toward MT sliding events and the remaining 94% of events retained a scissoring defect. In contrast, at 200 nM Eg5-Δtail-GFP about 45% of crosslinking events transitioned to form sliding events and 55% remained only scissoring events (Figure 4D). These data suggest the kinesin-5 tail-motor interface leads to slow motor motility, which is essential for concentrating Eg5 motors in the overlap zone. Increasing the number of motors in the overlap zone in the case of the Eg5-Δtail-GFP did not fully restore the zippering of two MTs into sliding zones, suggesting the tail-motor interface may be altering how motors undergo motility and interact with each of the two MTs.

### The tail domain is critical for slowing kinesin-5 motility within MT sliding zones

To understand the function of the tail-motor interface in kinesin-5 engagement within the MT sliding zones, we next studied how FL-Eg5-GFP or Eg5-Δtail-GFP motors interact with two MTs active sliding zones formed by non-fluorescent FL-Eg5 (Figure 4E). We carried out spiking experiments in which low concentrations of FL-Eg5-GFP or Eg5-Δtail-GFP were added to MT active MT sliding zones formed by FL-Eg5 (Figure 4E). MT sliding events were formed using 20 nM unlabeled FL-Eg5, and were spiked with either 1 nM FL-Eg5-GFP or 1 nM Eg5-Δtail-GFP to visualize single motor motility behavior along the anchored MT and within the MT sliding zones (Figure 4E; see materials and methods). We observe that FL-Eg5-GFP motors undergo motility along the anchored MT and enter into MT sliding zones. As they engage the free-MTs, their motility becomes slower and they remain plus-end directed along the anchored MT (Figure 4F,right panels; Video S5). In contrast, 1 nM Eg5-Δtail-GFP motors exhibit rapid plus-end directed motility along the anchored MT, outside the sliding zone and transition to bi-directional motility with extensive direction reversals, as they bind both MTs within overlap zones (Figure 4G, right panels; table 4; Video S5). We quantitated the pattern of motility using single motor tracking, and this revealed that motility velocity and the frequency of direction reversal along the anchored MT within MT sliding zones. FL-Eg5-GFP motors undergo slow plus-end directed along anchored MTs with very little direction reversal, and their motility velocity becomes slower within MT sliding zones (5-7 nm/sec) (Figure 4H). In contrast, Eg5-Δtail-GFP motors exhibit more rapid plus-end directed motility (10 nm/sec) along the anchored MT (Figure 4I). Upon entering MT sliding zones and interacting with both MTs, Eg5-Δtail-GFP motility remained rapid but the motors transitioned to bi-directional motility with frequent reversals (Figure 4I). We interpret these direction reversals as uncoordinated plus-end directed motility along either of the two anti-parallel MTs within sliding zones (Figure 4G, third panel from the left; Video S5). The Eg5-Δtail-GFP motility is roughly two-fold faster than FL-Eg5 GFP along single MTs and within MT sliding zones. The FL-Eg5-GFP motors remain attached within MT sliding zones for three-fold longer than Eg5-Δtail-GFP, which rapidly exited anti-parallel MT sliding zones. by moving bi-directionally towards the plus-end of either of the two sliding MTs. (Table 4; Figure 4F right panels; Figure 4I; Video S5). These data suggest that the Kinesin-5 tail regulates MT sliding rate by modulating a unique motor association with MTs and coupling between two MT bound ends of kinesin-5. Deletion of the tail leads rapid bi-directional motility with frequent reversals suggesting the two ends of kinesin-5 are both undergoing rapid uncoordinated motility cycles due to plus-end directed motility along both MTs within sliding zones.

### The tail domain is critical for kinesin-5 to generate MT sliding pushing forces

In order to understand how the tail regulates the kinesin-5 activity to slide apart antiparallel MTs, we next measured the forces generated by FL-Eg5-GFP or Eg5-Δtail-GFP on the free MT during sliding events using optical trapping combined with TIRF assays as previously described (Shimamoto et al., 2015)(Figure 5A). Here, we reconstituted anti-parallel pairs of MTs similar to those described above (Figure 4), but with an additional step of attaching polystyrene beads coated with nucleotide-free kinesin-1 motor domains to the ends of Rhodamine-labeled MTs (red). These bead-associated free-MTs were reconstituted into anti-parallel overlaps within flow chambers containing Hilyte-647-labeled MTs (purple) anchored to the slide surface and either FL-Eg5-GFP or Eg5-Δtail-GFP (green). These sliding MT pairs were then simultaneously imaged using TIRF microscopy while measuring the force exerted on the bead using an optical trap (Figure 5A). Under these conditions, 3-10 nM FL-Eg5-GFP produced robust MT sliding events (Figure 5B left panel; Video S6). In contrast, for 3-10 nM Eg5-Δtail-GFP, MT sliding was very rarely observed, and the majority of events involved free MTs undergoing ‘scissoring’ movements about a single point of the anchored MT, similar to our previous observations in TIRF sliding assays (Figure 4D). In order to recruit comparable amounts of Eg5-Δtail-GFP motors within the overlap regions and promote relative MT sliding, we needed to increase the amount of Eg5-Δtail-GFP in the chamber by 50-100-fold consistent with our previous results (Figure S4D; Figure 5B; Video S6). Under these conditions, regions of MT overlap exhibited increased recruitment of kinesin-5 molecules (Figure 5B; Video S6).

**Figure 5:**
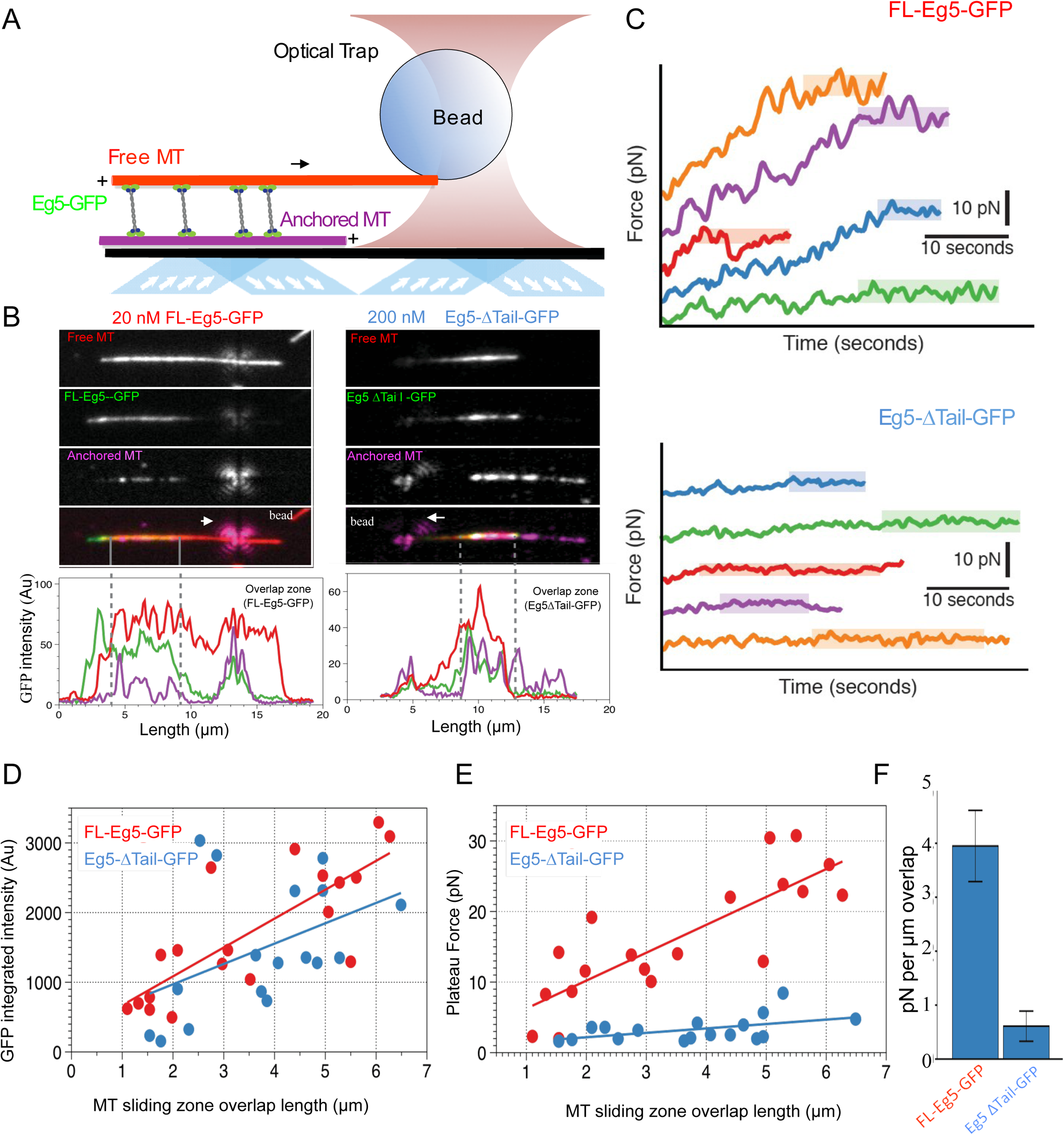
The kinesin-5 tail regulation is essential for generating pushing forces during MT sliding motility. A) Scheme for MT sliding and optical trapping to measure the MT sliding pushing forces. Hilyte 649 labeled and biotins labeled MTs (purple) were attached to glass surfaces via neutravidin-biotin. FL-Eg5-GFP motors (green) slide apart the Rhodamine-labeled MTs (red) while bound to a polystyrene bead (sphere) coated with kinesin-1 rigor mutant protein, which becomes locked into the optical trap to measure forces. B) Example images of MT sliding events mediated by FL-Eg5-GFP (left) and Eg5-Δtail-GFP (right). Top, free MT shown in gray scale. Second, Eg5-GFP intensity in the overlap zone. Third, attached MT. The polystyrene bead can be seen attached to the free MT. Lower panel, fluorescence intensity for each of the channels above showing the identification of the overlap zone length and total GFP intensity above background. C) Example optical trapping force profiles generated for sliding events by FL-Eg5-GFP (top) and Eg5-Δtail-GFP (bottom). Top panel, FL-Eg5-GFP MT sliding events generate and build up forces that then plateau (highlighted level). Lower panel, Eg5-Δtail-GFP MT sliding events generate very weak forces, which plateau at lower values. D) Scaled comparison for overlap zone length (µm) plotted in relation to the overall GFP intensity for each sliding event. FL-Eg5-GFP is shown in red while Eg5-Δtail-GFP is shown in blue. Note there is generally little discernable statistical difference between the slopes of these comparisons. E) Scaled comparison of the plateau forces generated (pN) in relation to the size of the overlap zone for each MT sliding event. FL-Eg5-GFP is shown in red while Eg5-Δtail-GFP is shown in blue. Note the slope of the FL-Eg5-GFP force to length comparison is steeper than the Eg5-Δtail-GFP force to length comparison F) Forces generated per µm of overlap length representing the slope of linear comparison in E.

Once sliding of the free MT was observed, the Rhodamine-labeled MT-bound polystyrene bead was optically trapped and the magnitude of the pushing force generated by MT sliding was measured (Figure 5C). FL-Eg5-GFP mediated MT sliding exhibited a steady increase in the force produced until a maximum ‘plateau’ force was reached, frequently resulting in many 10s of pN of force across the MT sliding pair. This behavior was quite similar in both time-scale and force magnitude to reports of *Xenopus* Eg5 pushing forces measured in a similar assay (Shimamoto et al, 2015). In contrast, Eg5-Δtail-GFP-mediated MT pairs exhibited short excursions of force increase, reaching significantly lower ‘plateau’ values and lower overall total forces generated (Figure 5C).

For each individual MT sliding event examined, we calculated the integrated intensity of GFP signal within the overlap region as defined by the distance between MT plus-ends (dashed gray lines, Figure 5B). The integrated intensity served as a proportional readout to the amount of motors located within the overlap region. By comparing many MT sliding zones, we were able to determine that an approximately linear relationship emerged between the length of relative MT overlap zone length and the number of motors within this region (Figure 5D). Furthermore, at the concentrations of FL-Eg5-GFP and Eg5-Δtail-GFP used in these assays, a comparable relationship was seen for both constructs, suggesting that similar numbers of motors were engaged within overlaps of similar size (Figure 5D).

We also examined the relationship between the magnitude of the force plateau reached in each individual MT sliding event and the length of the MT overlap zone during force generation. For both Eg5-GFP and Eg5-Δtail-GFP a nearly linear increase in plateau force relative to MT overlap zone length was observed (Figure 5E). However, the slopes of these relationships differed significantly between the two constructs. Here the slopes represent the plateau pushing force generated per μm length of sliding zone: the Eg5-Δtail-GFP generates about 0.6 ± 0.3 pN per μm while the FL-Eg5-GFP generated about 4.0 ± 0.6 pN per μm, indicating about seven-fold difference in MT sliding forces (Figure 5F). Therefore, the pushing forces produced by sliding anti-parallel MT bundles by ensembles of Eg5-Δtail-GFP are roughly seven-fold lower than that those generated by MT sliding bundle ensembles of comparable numbers of FL-Eg5-GFP motors. Together, these data indicate that the tail domain is critical for regulating the homotetrameric kinesin-5 motor to produce pushing forces during MT sliding events.

### The tail domain is critical for Eg5 localization to metaphase and anaphase mitotic spindles

We next determined the role of the kinesin-5 tail domain in motor localization of Eg5 in metaphase and anaphase in HeLa cells. Cells were transfected with human GFP-α-tubulin to visualize MTs, and either FL-Eg5-mCherry (FL-Eg5-mCh), Eg5-Δtail-mCherry (Eg5-Δtail-mCh), or mCherry alone (mCh). Expression of each construct in HeLa cells was assessed by western blot (Figure 6A). In fixed, metaphase cells, FL-Eg5-mCh localized to spindle MTs, while in contrast localization of Eg5-Δtail-mCh was more diffuse, with increased cytoplasmic signal (Figure 6B). This difference in localization was quantified as the ratio of mCh signal localized to the mitotic spindle and mCh signal in the cytoplasm. In cells with comparable mCh-construct expression levels (Figure 6C, left), this spindle to cytoplasm ratio was high for FL-Eg5-mCh (2.39 ± 0.08, mean ± SEM) and significantly reduced for Eg5-Δtail-mCh (1.37 ± 0.02, p<0.0001, ANOVA with Tukey’s multiple comparisons test), indicating that deletion of the tail causes a defect in localization to metaphase spindle MTs (Figure 6C, right). Treatment with the compound BRD-9876, which locks Eg5 in a nucleotide-free-like state (Chen et al., 2017), only partially rescues the localization of Eg5-Δtail-mCh to metaphase spindle MTs, potentially due to defects in both the on and off-rates (Figure 6D, 6E).

**Figure 6:**
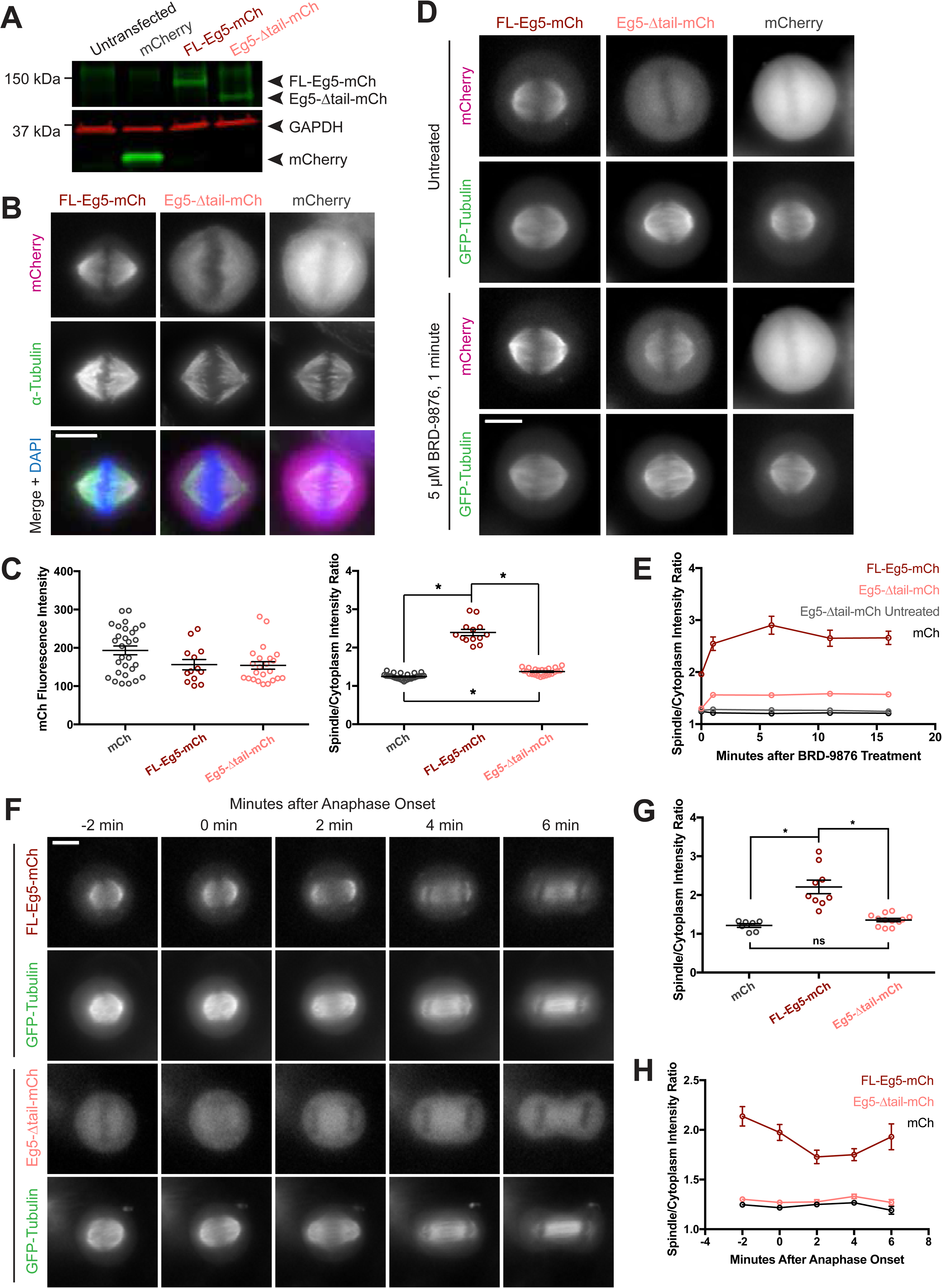
Deletion of the kinesin-5 tail domain disrupts mitotic spindle localization in metaphase and anaphase. A) Western blot for mCherry (mCh, green) and GAPDH (red) indicating the expression of FL-Eg5-mCherry (FL-Eg5-mCh, green) and Eg5-Δtail-mCherry (Eg5-Δtail-mCh, green) in HeLa cells. Results are representative of three independent experiments. B) Localization of mCh, FL-Eg5-mCh, and Eg5-Δtail-mCh in HeLa cells arrested in metaphase via treatment with MG-132. FL-Eg5-mCh localizes to spindle MTs. Tail deletion disrupts localization, and Eg5-Δtail-mCh signal is distributed between spindle MTs and the cytoplasm. Scale bar 10 μm. C) Left panel, mCh fluorescence intensities of single cells used for quantification of localization to spindle MTs (n = 13-29 cells per transfection condition, three independent experiments). Right panel, the ratio of mCh fluorescence signal on the spindle to signal in the cytoplasm is significantly lower in fixed metaphase cells expressing Eg5-Δtail-mCh compared to FL-Eg5-mCh, indicating reduced localization of Eg5-Δtail-mCh to spindle MTs (n = 13-29 cells per transfection condition, three independent experiments, * indicates p < 0.01, ANOVA with Tukey’s post hoc test). D) Treatment of live HeLa cells expressing Eg5-mCh constructs and GFP-Tubulin with the Eg5 rigor inhibitor BRD-9876 results in a rapid (< 1 minute) increase in FL-Eg5-mCh signal on the spindle. Inhibitor treatment increases, but does not fully rescue, localization of Eg5-Δtail-mCh to the spindle (n = 7-13 cells per transfection condition, three independent experiments). E) The ratio of mCh fluorescence signal on the spindle to signal in the cytoplasm rapidly increases after treatment with BRD-9876 in cells expressing FL-Eg5-mCh or Eg5-Δtail-mCh. The spindle to cytoplasm intensity ratio of Eg5-Δtail-mCh expressing cells never reaches that of cells expressing FL-Eg5-mCh, indicating only partial rescue of motor localization with rigor inhibitor treatment. BRD-9876 treatment does not alter the ratio of mCh control cells. F) Deletion of the tail domain disrupts localization of Eg5 to the spindle in anaphase. Paired rows of images demonstrate the localization of FL-Eg5-mCh and Eg5-Δtail-mCh as HeLa cells expressing GFP-tubulin transition from metaphase to anaphase. FL-Eg5-mCh signal is observed at the spindle throughout the metaphase to anaphase transition and the motor localizes to the midzone after anaphase onset (see 4-6 min panels). Increased cytoplasmic and reduced spindle signal is observed in cells expressing Eg5-Δtail-mCh throughout the metaphase to anaphase transition. G) The ratio of mCh fluorescence signal on the spindle to signal in the cytoplasm was measured six minutes after anaphase onset. As in metaphase cells, localization of Eg5-Δtail-mCh to the spindle was significantly reduced compared to FL-Eg5-mCh (n = 7-12 cells per transfection condition, three independent experiments, * indicates p < 0.01, ANOVA with Tukey’s post hoc test). H) The spindle to cytoplasm intensity ratio of cells expressing Eg5-Δtail-mCh is lower than that of cells expressing FL-Eg5-mCh throughout the metaphase to anaphase transition, indicating a persistent localization defect caused by deletion of the tail domain (n = 7-12 cells per transfection condition, three independent experiments).

Live cell imaging was used to assess whether this tail deletion localization defect was present in anaphase as well as metaphase mitotic spindles. Measurement of spindle to cytoplasm mCh intensity ratios in anaphase showed a similar reduction, indicating decreased localization to spindle MTs, for Eg5-Δtail-mCh compared to FL-Eg5-mCh as seen in metaphase cells (ratios 1.36 ± 0.04 and 2.21 ± 0.17, mean ± SEM, respectively, p < 0.0001, ANOVA with Tukey’s multiple comparisons test) (Figure 6F, 6G). As HeLa cells progress from metaphase to anaphase, the deletion of the tail consistently reduces the localization of Eg5 to spindle MTs (Figure 6F, 6H). These data support a critical role for the tail domain in kinesin-5 mitotic spindle MT localization.

## Discussion

Kinesin-5 motors share a conserved anti-parallel MT sliding activity, which is essential for the assembly and elongation of mitotic spindles. This conserved activity allows these motors to mobilize both their MT tracks simultaneously as cargos being transported. Here, we show that a conserved tail-to-motor interaction at the two ends of the kinesin-5 homotetramer is responsible for promoting the unique kinesin-5 MT sliding motility (Figure 7A-B). The regulation is most clearly observed at low ionic strength conditions, where the tail exhibits the highest affinity to the motor domain and the highest level of MT activated motor ATP hydrolysis regulation leading to a slow and high affinity MT catalysis cycle (Figure 7B). Structurally, the tail domain binds the motor domain in either the nucleotide-free or ADP MT-bound states by engaging the N-terminal subdomain at α0-helix hairpin (Figure 7C). This element of the kinesin-5 motor domain rotates upward from the MT lattice in the nucleotide free state compared to its downward position in the ATP state (Figure 2; Figure 7C). We observe processive FL-Eg5-GFP motility along individual MTs and this was previously not observed likely due to lower than physiological pH (pH 6.8) *in vitro* conditions utilized in previous studies (Kapitein et al., 2008). The tail-motor interaction down-regulates MT activated ATP hydrolysis resulting in very slow motility for FL-Eg5-GFP at 25 mM KCl along individual MTs and within MT sliding zones (Figure 3–4). The tail-motor interaction is overcome partially by increasing the solution ionic strength (50-100 mM KCl), which weakens tail-motor interface partially, relieving the suppression of ATP hydrolysis and leading to increased motility velocities. The tail also enhances the clustering of kinesin-5 homotetramers into oligomeric assemblies. At higher ionic strength (50 mM KCl), the MT sliding velocity directly correlates with an increase in the motility velocity of FL-Eg5-GFP motors within MT sliding zones. In contrast, Eg5-Δtail-GFP motors exhibit no suppression in motor velocity at 25 mM KCl, which remained mostly constant at higher ionic strengths (50-100 mM KCl), and exhibited no clustering behavior. The Eg5-Δtail-GFP velocity remained consistently 20% higher than Eg5-FL-GFP at 50 and 100 mM KCl, suggesting some tail regulation remains in place in those conditions. However, weakening the tail to motor interface enhances the higher-order clustering among FL-Eg5-GFP homotetramers at 50-100 mM KCl, suggesting a transition of tail from intra to inter motor homotetramer interactions maybe responsible for clustering (Figure 7A, broken lines). The tail domain also enhances processive motility run lengths for FL-Eg5-GFP at higher ionic strength a feature that was 45% decreased for Eg5-Δtail-GFP (Figure S3B). These data are fairly consistent with previous observations for full-length and tail-deleted *Xenopus* Eg5 (Weinger et al., 2011).

**Figure 7:**
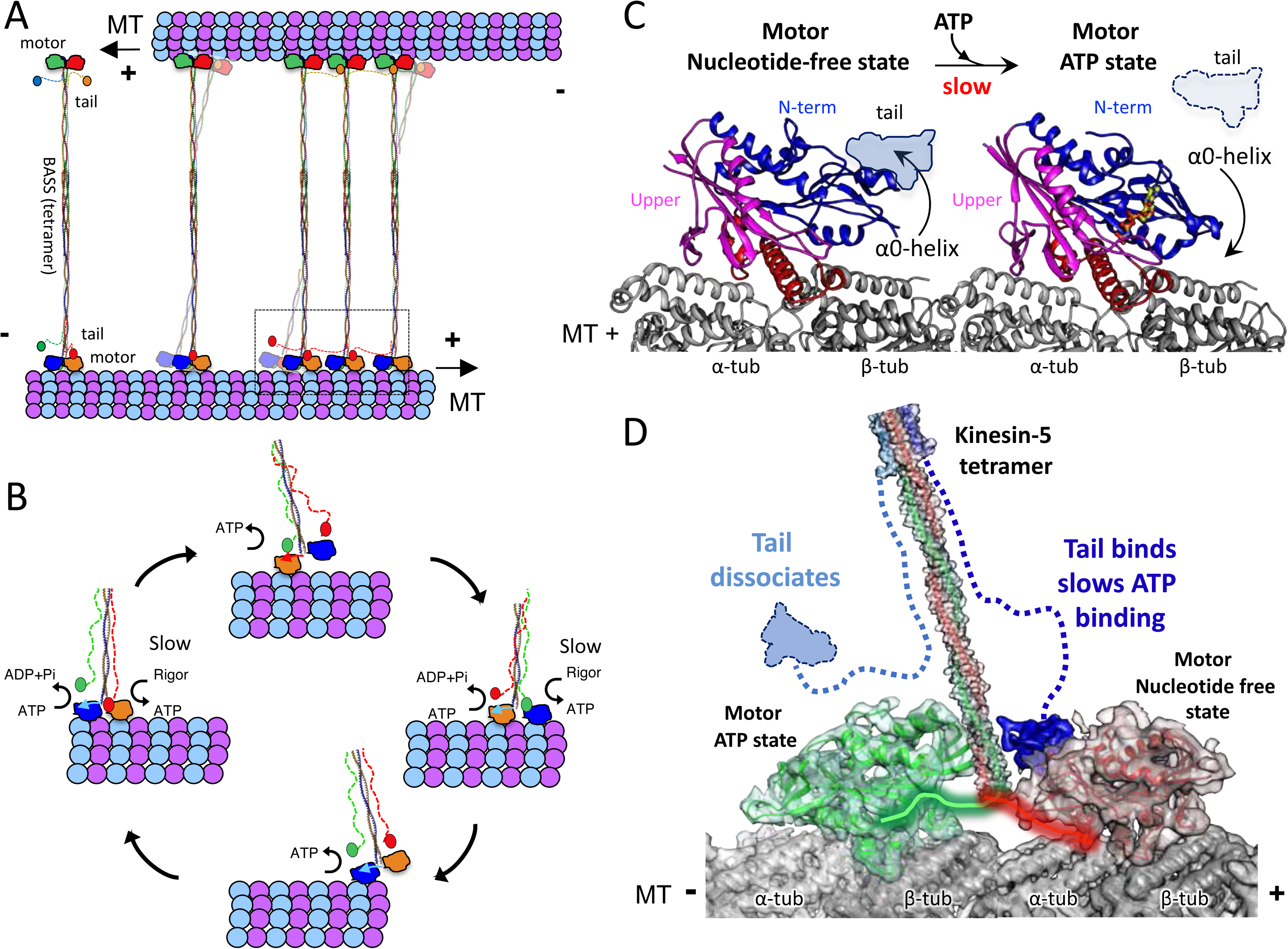
A mechanistic model for kinesin-5 tail-motor domain interaction during walking motility and the critical role for this interaction during MT sliding activity. A) Model for kinesin-5 homotetramers with their motor and tail domains at each end of the bipolar minifilament (60 nm). The motors and tail domains at each end may form assemblies where tail domain of one homo-tetramer make contact with motors of a second tetramer to form clusters of two to four motors B) The role of the tail in regulating the kinesin-5 hand-over-hand motility cycle by slowing ATP binding of the lead motor leading to slow hand over-hand motility and prevalence of the dual bound state at each end of kinesin-5. C) The conformations of the kinesin-5 motor subdomains in the process of ATP binding. Left, The N-terminal subdomain (blue) is in upward state with the helix-0 to engage the tail domain and wedging the nucleotide binding pocket open. Right, the N-terminal subdomain moves downward enclosing on the bound bound ATP, leading the helix-0 to move downward and disengage from the tail domain. D) Synthesized view of dual dimeric motor bound state of the kinesin-5 motor end. This state was synthesized based on the cryo-EM maps and, in vitro reconstitution, biochemical and kinetic studies described here. The tail makes contact with the motor domain only in the nucleotide-free state but dissociates when the motor is in the ATP state. The lead motor is bound to the tail while the trailing motor dissociates from the tail domain.

Our studies suggest that tail-to motor interface occurs at both ends of the kinesin-5 homotetramer and is critical for coordinating their motility activities required for promoting MT sliding motility. We show that human FL-Eg5-GFP motors capture antiparallel MTs and zipper them into sliding MT bundles (Figure 4; Figure S4). Loss of the tail-to-motor interface in Eg5-Δtail-GFP motors has no effect on the initial crosslinking of two MTs at or near MT-plus-ends, but leads to severe defect in zippering the two MTs into sliding bundles (Figure 4). Within active MT sliding zones, we show that the Eg5-Δtail-GFP motors are persistently bi-directional, by undergoing motility along either antiparallel MTs likely due to loss of coordination between the two bipolar kinesin-5 ends (Figure 4F, left panel). In contrast, FL-Eg5-GFP undergoes smooth, slow and unidirectional motility towards the MT plus-end of the anchored MT. These FL-Eg5-GFP motors generate seven-fold higher pushing forces compared to Eg5-Δtail-GFP motors while sliding apart antiparallel MTs. The productive forces generated by FL-Eg5-GFP result from the kinesin-5 tail to motor interface modulating motility at both MTs within the sliding zone. The tail enhances the FL-Eg5-GFP motor MT affinity leading to tightly engaged and slow moving motors onto both MTs within sliding zone.

### A modified model for kinesin-5 tail regulated hand-over-hand motility and its essential role in antiparallel MT sliding activity

Our studies suggest that tail domain regulates processive hand-over-hand stepping during kinesin-5 motility (Figure 7B-D)(Vale and Milligan, 2000). Well-studied MT plus-end directed kinesin motors (i.e. Kinesin-1) undergo hand-over-hand walking motility along MT protofilaments utilizing dimeric motor domains coupled by their neck linkers and neck coil-coils. During hand-over-hand motility, one motor domain of a dimeric kinesin binds the MT lattice in a trailing position, while the other motor domain occupies a lead position, 8-nm apart along two consecutive αβ-tubulins within MT protofilaments (Figure 7D). The trailing motor forms an ATP-like state with its neck-linker docked, while the lead motor initially binds weakly leading to ADP dissociation nucleotide-free state (Figure 7D). The hand over hand cycle of dimeric kinesin-1 motors result in 8-nm steps toward the MT plus-ends (Vale and Milligan, 2000). Our studies suggest a new form of regulation for hand-over-hand motility, in which the kinesin-5 tail domain down-regulates ATP binding in the lead motor domain and stabilizes its ADP or nucleotide-free state (Figure 7C-D). The kinesin-5 tail binds the α0-helix within the motor domain, stabilizing its N-terminal subdomain in an upward conformation stabilizing an open active site, and slowing its ATP binding capacity (Figure 1–3; Figure 7C-D). This model is consistent with most of our data and indicates that the tail-motor regulation is the source of the conserved kinesin-5 pushing force generating activity that is essential for anti-parallel MT sliding. In yeast kinesin-5 orthologs, such as cin8, this interactions may further regulate the reversal of direction from minus-end directed motility along single MTs to plus-end directed motility within sliding zones as deletion of the tail domain leads to slow plus-end directed motility as tail deletion in Cin8 interferes with directionality reversal (Duselder et al., 2015).

This regulation model suggests that as kinesin-5 homotetramers undergo hand-over hand motility along two anti-parallel MTs within sliding zones, the two sets of tail domains emanating from oppositely folded subunits may bind the lead motor domains in either the ADP or nucleotide-free states stabilizing a double bound motor state along each MT. The tail-motor interface likely stiffens both two kinesin-5 interactions with both sliding MTs, and this likely improves force transmission between their two bipolar ends (Figure 5)(Figure 7B-D). Our study suggest that the tail-motor interaction increases the time each kinesin-5 dimeric end spends in the dual-bound motor states. In contrast to FL-Eg5-GFP, Eg5-Δtail-GFP motors are unable to effectively engage both sliding MTs due to the rapid motility, exhibit reversal in the directional motility by moving on both sliding MTs asynchronously, and exhibit a poor capacity for generating MT sliding pushing forces (Figure 4–5). The tail-motor interfaces effectively maintain the high affinity state for the lead motors at both bipolar kinesin-5 homotetramer ends, leading to efficient force transmission across its central minifilament (Figure 7C-D). Deletion of the kinesin-5 tail leads to rapid hand-over-hand stepping motility by either of the kinesin-5 ends, but severely decreased sliding forces due to defects in communication between the two kinesin-5 homotetramer ends.

Our data indicate that this interaction may be responsible for clustering multiple homotetramers into larger complexes (Figure 7A). Clusters of up to four Kinesin-5 motors were observed in the yeast ortholog, Cin8, where it induces motility reversal of direction forms the site for capturing free MTs to promote MT sliding (Shapira et al., 2017). In combination with data from our study, we suggest that clustering of Kinesin-5 motors is a general feature and it may be critical for coordinating multiple motor motility cycles within MT sliding zones (Figure 7A). Motor clusters consisting of up to two to four homotetramers move slowly in synchrony. Such clusters do not form in Eg5-Δtail motors. Another possibility may involve cooperativity among multiple motors in a traffic jam due to slow stepping, which was also previously suggested for cut7, the *S. pombe* ortholog (Britto et al., 2016).

### Kinesin-5 Tail to motor domain regulation is likely modulated by mitotic kinases is critical for mitotic functions

The kinesin-5 tail domain is essential for mitotic spindle assembly and elongation functions. Our studies confirm the defect in Eg5-Δtail-GFP may relate to the rapid cycling on and off the mitotic spindle MTs; trapping Eg5-Δtail-GFP motor in the nucleotide-free state using chemical strategies dramatically enhances their MT spindle localization (Figure 6). A recent study identified Eg5 mutations in culture cells leading to its chemical resistance to Eg5 inhibitors. Among these is a mutant cell strains possessing an Eg5 mutant trapped in the nucleotide-free state which can form near normal bipolar metaphase spindle that restores mitotic organization in the presence of the compounds (Sturgill et al., 2016). These data are consistent with the critical importance of the nucleotide-free state in the force generation cycle of kinesin-5, which can be stabilized by the tail-motor interface.

The tail-motor interface is essential for the stable localization of kinesin-5 motors to mitotic spindle MTs. The Kinesin-5 tail domain contains the conserved BimC box, a mitotic CyclinB dependent kinase phosphorylation consensus site (Blangy et al., 1995; Sharp et al., 1999). This motif is conserved across kinesin-5 orthologs and its mitotic phosphorylation was shown to be critical for the accumulation of kinesin-5 motors in the mitotic spindle midzones as it promotes promote spindle elongation in anaphase (Sharp et al., 1999). It remains unclear how this BimC box phosphorylation influences the kinesin-5 tail to motor regulation, which we describe here. However, it is potentially possible that the mitotic phosphorylation may further enhance the tail - motor interface and further slowing MT-based motility. This may represent a more effective form of kinesin-5 regulation which may lead to more efficient MT sliding function during anaphase. Future studies of the phosphorylation modified Kinesin-5 motors at the BimC box motifs may help lay out the detailed effect on regulating kinesin-5 motility.

### Concluding Remarks

We reveal a new mechanism for interaction between the kinesin-5 tail and motor domain during motility along MTs. The tail binds the motor domain and stabilizes it in the high affinity MT bound nucleotide-free state. This regulation is essential for slowing kinesin-5 walking motility and is central for transforming walking motility to slide apart MTs a feature that is essential for the mitotic function of kinesin-5

## Materials and Methods

### Generation of Constructs for various studies

For *in vitro* motility studies, we generated constructs for expression insect cells full length coding regions for Human Eg5 (KIF11)(FL-Eg5: residues 1-1056) and C-terminally truncated Eg5 (Eg5-Δtail; residues 1-920) were inserted in pFastbac1 vector in frame of a C-terminal StrepII tag or fused to monomeric superfolder-GFP(msf-GFP) and a C-terminal Strep-II tag.

For *in vivo* imaging studies, A pcDNA-3.1 plasmid containing Eg5 full-length fused to mCherry (Eg5 FL-mCh) inserted between the EcoR1 and Not1 sites was used as a template. Plasmid pAT4206, encoding Eg5-Δtail, which encoded residues 1-912, fused to mCherry (Eg5 ΔTail-mCh), was generated using the Phusion Site-Directed Mutagenesis Kit (Thermo Scientific). These clone contained silent mutations for siRNA-resistance: T2124C, C2130T, G2133T, and G2136A. The forward and reverse primer sequences used to generate Eg5 ΔTail-mCh were 5’-GGAGCGCCAATGGTGAGCAA-3’ and 5’-AAAGCAATTAAGCTTAGTCAAACCAATTTT-3’, respectively.

For Kinetic and structural studies, coding regions for isolated Dm KLP61F and Human Eg5 motor-neck linker constructs (KLP61F motor residues 1-369; Eg5 motor residues 1-360) and the tail domains for these proteins (KLP61F tail residues 906-1016 and Eg5 residues 920-1056) and KLP61F motor-tail fusion (residues 1-369 fused to 906-1016) were inserted in frame with histidine-tag in pET v2 vector (macrolab UC-berkeley)

### Protein expression and Purification

For expression FL-Eg5-GFP, FL-Eg5 and Eg5-Δtail-GFP was carried out using baculovirus expression system. Briefly 200 to 500 mL of Spodoptera frugiperda (Sf9) cells, were infected with third passaged virus for each construct and expressed for 60-72 hours. Virus-Infected Sf9 Cells were centrifuged at 1500 rpm and then washed and incubated with lysis buffer (50 mM HEPES 300 mM KCl, 10 mM beta-meractoptoethanol, 1 mM MgCl2, 0.2 mM ATP) in the presence of 0.5% Triton X100. Cells were lysed by extrusion using a dounce homogenizer and then were centrifuged at 40,000 rpm using Ti45 rotor in ultracentrifuge (Beckman). The lysate was passaged on a Streptactin XT resin (IBA lifesciences) equilibrated with lysis buffer, washed extensively and then eluted with lysis buffer supplemented with 100 mM D-Biotin. Purified Eg5 proteins were concentrated and loaded onto Superose-6 (10/300) column (GE Health Care) equilibrated onto AKTA system and eluted in 0.5 mL fractions (Figure S3A-B). Fractions were evaluated using SDS-PAGE for protein quality and Eg5 protein containing fractions were either used immediately for motility experiments or were snap frozen with 15% glycerol in lysis buffer using liquid nitrogen. We did not observe any difference in the FL-Eg5-GFP, FL-Eg5 and Eg5-Δtail-GFP activities in the frozen or freshly prepared settings (Figure S3A-B).

For kinetic and structural cryo-EM studies, KLP61F motor, tail and motor-tail constructs and Eg5 motor and tail constructs were expressed using T7 expression in SoluBL21 cells. Briefly, cells were grown at 37°C and induced for expression using 0.5 mM Isopropyl thio-beta-glucoside overnight at 19 °C. Cells were centrifuged and lysed in Lysis buffer using a microfluidizer. Proteins were purified using IDA-nickel affinity chromatography (Macherey Nagel), washed extensively and then eluted with lysis buffer supplemented with 200 mM Imidazole. The purified fractions were loaded onto a Superdex 200 (16/6) size exclusion column equilibrated with lysis buffer at 1 mL fractions. SDS-PAGE was used to evaluate fractions and concentrated pure proteins were incubated with 15% glycerol before freezing in liquid nitrogen.

### MT activated ATP hydrolysis and MT-co-sedimentation assays

MT-activated ATPase activity for KLP61F motor, tail, motor-tail fusion and Eg5-motor and tail constructs were carried out as previously described. We determined in 50 mM potassium acetate or 20-80 mM KCl, 25 mM HEPES, 5 mM magnesium acetate, 1 mM EGTA, pH 7.5 buffer by measuring phosphate production in the presence of a minimum of a 5-fold molar excess range of MT concentrations, using a commercially-available kit (EnzChek, Molecular Probes) at 20°C.

To measure MT co-sedimentation activity for KLP61F and Eg5 motor and tail constructs, we prepared 5 mg/ml MTs in 5% DMSO by polymerization at 37°C and then stabilized with Paclitaxel (sigma), MT co-sedimentation assays were carried out by mixing 0 to 2.5 mg/ml KLP61F motor with or without tail domain, or motor-tail in 50 mM HEPES pH 7.5, 5mM MgCl2, 1% glycerol, 25 mM or 75 mM KCl in the presence of 2 mM AMPPNP, 2 mM ADP, 2 mM ADP.AlF4 or 10U Apyrase to mimic the nucleotide-free state. Mixtures of motors, tail proteins bound to MTs in these conditions were incubated at 25°C for 20 min and then centrifuged at 18k for 25 min at 25°C. The supernatant was then removed and mixed with SDS sample buffer. The pellets were resuspended with SDS sample buffer. Equal amounts of each supernatant and pellet were analyzed by SDS PAGE and presented. Intensity quantitation was carried out using Biorad Image Lab software (Biorad).

### Sample preparation of motor and tail decorated MTs for Cryo-EM

MTs were prepared by polymerizing 5 mg/ml tubulin (Cytoskeleton, Denver, CO) in BRB80 buffer (80 mM PIPES, pH 6.8, 1 mM EGTA, 4 mM MgCl2, 2 mM GTP, 9% dimethyl sulfoxide) for 30 min at 36° C. Paclitaxel was added at 250 μM before further incubation of 30 min at 36°C. The polymerized MTs were then incubated at room temperature for several hours or overnight before use. For grid preparations, KLP61F motor and KLP61F tail proteins were mixed in 0.5% binding buffer (25mM HEPES, 35 mM potassium acetate plus 2mM ADP to a final concentration of 0.1mg/mL and 0.04mg/mL respectively). For grid preparation of motor alone KLP61F motor (10mg/mL) was diluted in binding buffer (50mM HEPES, 70 mM potassium acetate plus 2mM AMPNP). All samples were prepared on 1.2/1.3 400-mesh grids (Electron Microscopy Services). Grids were glow-discharged before sample application. The cryo-samples were prepared using a manual plunger, which was placed in a homemade humidity chamber that varied between 80 and 90% relative humidity. A 4-μl amount of the MTs at ~0.5 μM in 80 mM PIPES, pH 6.8, 4 mM MgCl2, and 1 mM EGTA supplemented with 20 μM Paclitaxel was allowed to absorb for 2 min, and then 4 μl of the KLP61F motor and tail domains were added to the grid. After a short incubation of 2 min, 0.5 μL of Apyrase (25 units) was added and after a short incubation of 3 min the sample was blotted (from the back side of the grid) and plunged into liquid ethane. This procedure was repeated for the motor alone preparations without the addition of the apyrase step.

### Cryo-EM Image analysis, structure determination and model building

Images of frozen-hydrated KLP61F motor decorated MTs in the AMPPNP state or KLP61F motor and tail decorated MTs in the nucleotide-free state (see Supplemental Table1) were collected on a Titan Krios (FEI, Hillsboro, OR) operating at 300 keV equipped with a K2 Summit direct electron detector (Gatan, Pleasanton, CA). The data were acquired using the Leginon automated data acquisition (Suloway et al., 2005). Image processing was performed within the Appion processing environment (Lander et al., 2009). Movies were collected at a nominal magnification of 22500× with a physical pixel size of 1.31 Å/pixel. Movies were acquired using a dose rate of ~7.96 and 8.3 electrons/pixel/second over 8.25 seconds yielding a cumulative dose of ~38 and 40 electrons/Å2 (respectively). The MotionCor frame alignment program (Hirschi et al., 2017; Li et al., 2013) was used to motion-correct. Aligned images were used for CTF determination using CTFFIND4 (Rohou and Grigorieff, 2015) and only micrographs yielding CC estimates better than 0.5 at 4 Å resolution were kept. MT segments were manually selected, and overlapping segments were extracted with a spacing of 80 Å along the filament. Binned boxed segments (2.62Å/pixel, 192 pixel box size) were then subjected to reference-free 2D classification using multivariate statistical analysis (MSA) and multi-reference alignment (MRA) (Hirschi et al., 2017; Ogura et al., 2003). Particles in classes that did not clearly show an 80Å layer line were excluded from further processing.

For cryo-EM reconstruction, undecorated 13,14- and 15-protofilament MT densities (Sui and Downing, 2010) were used as initial models for all preliminary reconstructions. We used the IHRSR procedure (Egelman, 2007) for multi-model projection matching of MT specimens with various numbers of protofilaments (Alushin et al., 2014), using libraries from the EMAN2 image processing package (Tang et al., 2007). After each round of projection matching, an asymmetric back-projection is generated of aligned segments, and the helical parameters (rise and twist) describing the monomeric tubulin lattice are calculated. These helical parameters are used to generate and average 13, 14 and 15 symmetry-related copies of the asymmetric reconstruction, and the resulting models were used for projection matching during the next round of refinement. The number of particles falling into the different helical families varied. Helical families that had enough segments were further refined. Final refinement of MT segment alignment parameters was performed in FREALIGN (Grigorieff, 2007) without further refinement of helical parameters. FSC curves were used to estimate the resolution of each reconstruction, using a cutoff of 0.143. To better display the high-resolution features, we applied a B-factor of 200 Å, using the program bfactor (http://grigoriefflab.janelia.org).

In order to enhance the total mass of the mobile tail density and improve the resolution of the MT decorated with KLP61F motor and tail in the nucleotide-free state, an additional round of “MT-patch refinement” processing was performed enhance conformational homogeneity. The same motion-corrected micrographs and boxes were used, but defocus parameters were re-estimated using GCTF (Zhang, 2016). MTs were then sorted into 13, 14, and 15 protofilament MTs using reference alignment as previously described (Shang et al., 2014). Of the 29,274 starting particles, 19,128 MT particles corresponding to the 14 protofilament symmetry were selected for further processing. MTs were then refined using RELION helical processing (He and Scheres, 2017). Initially, the asymmetric unit was defined as one full 82Å repeat (consisting of 13 tubulin dimers), using an initial estimate of zero for the helical twist. Local symmetry searches were performed during the refinement to optimize these parameters. Following refinement, the particle coordinates were smoothed as previously described (Huehn et al., 2018). After MT refinement, an additional protofilament refinement step was performed in an attempt to increase resolution of the final volume by correcting for distortions in the MT lattice. To do this, a wedge mask is applied to the final MT volume, resulting in a MT missing a single protofilament. This volume was then rotated and subtracted thirteen times from each experimental image to generate a stack of protofilament particles, with one particle for every tubulin dimer in the imaged filament. In this case, 267,792 protofilament particles were obtained from the original stack of 19,128 MT particles. Protofilament particles alignment parameters were initialized using euler angles derived from the MT refinement step and subjected to further, local refinement using RELION. The final resolution was computed using the RELION post-processing module with a soft-edged mask. A more detailed description of the protofilament refinement method is currently being published (Debs et al. manuscript in preparation). In order to more accurately estimate the resolution of each region of the reconstructed density, a local resolution calculation was performed using the “blocres” function in the Bsoft processing package (Heymann and Belnap, 2007). This analysis revealed that the majority of the tubulin density is in the range of 3.5 -4.5Å, while the kinesin portion ranges from 5-6Å resolution and tail density is around 8 Å resolution (Figure S2A-C). Model building was performed using the programs Coot and UCSF chimera using the kinesin-5 structural model (Turner et al., 2001). The model was adjusted using the secondary structure elements in the density maps. The maps were compared using the programs UCSF chimera and Coot to determine the transitions of various elements.

### Reconstitution of Eg5 motility along single MTs

Kinesin-5 MT activated motility was reconstituted as follows: Flow chambers were assembled from N 1.5 glass coverslips (0.16 mm thick; Ted Pella) that were cleaned with the Piranha protocol and functionalized with 2 mg/mL PEG-2000-silane containing 2 μg/mL biotin-PEG-3400-silane (Laysan Bio) suspended in 80% at pH 1 (Henty-Ridilla et al., 2016). After the flow chamber was assembled, 0.1 mg/mL NeutrAvidin (Thermofisher) was used to functionalize surfaces. Biotin and Alexa-Fluor-633-labeled porcine tubulin were generated in the laboratory as described (Al-Bassam, 2014) and were polymerized using the non-hydrolyzable GTP analog guanosine-5’-[(α,β)-methyleno] triphosphate (termed GMPCPP; Jena Biosciences) or using the MT stabilizing drug, Paclitaxel (sigma). These MTs (100-200 μg/mL in BRB-80: 80 mM PIPES, 1 mM MgCl_2_ and 1 mM ETGA; pH 6.8, 1% glycerol, 0.5 % pluronic-F127, 0.3 mg/ml casein, 3 mM BME, 4 mM ATP-MgCl2) were flowed into chambers and attached to glass via biotin-neutravidin linkage. Flow chambers were then extensively washed with imaging buffer (25 mM HEPES, 25-100 mM KCl, pH 7.5, 10 mM beta-mercatopethanol; 1% glycerol, 0.5 % pluronic, 0.3 mg/ml casein, 3 mM BME, 4 mM ATP-MgCl2). Kinesin-5 MT-activated motility was reconstituted at 25 °C by injecting 1-20 nM FL-Eg5-GFP combined with a photobleach-correction mixture into flow chambers (Telley et al., 2011). Movies were captured in TIRF mode using a Nikon Eclipse Ti microscope using 1.5 Na objective and an Andor IXon3 EM-CCD operating with three (488 nm, 560 nm and 640 nm) emission filters using alternating filter wheel in 2-s increments operated using elements software (Nikon)

### Reconstitution of Eg5 MT sliding motility

To study MT sliding activities *in vitro*, flow chambers were prepared as described above and either Paclitaxel or GMPCPP stabilized AlexaF-633 and biotin labeled MTs were anchored along their surface via Biotin-Neutravidin linkage. A mixture of 1-20 nM FL-Eg5-GFP or 20-200 nM Eg5-Δtail-GFP were mixed with 100-200 μg/ml AlexaF-560 labeled MTs and injected into these flow chambers, after being equilibrated with imaging buffer. Imaging was initiated as described above almost immediately and areas of MT sliding events were identified through search. At 3-20 nM FL-Eg5-GFP robust free MT crosslinking (yellow) was observed, followed by zippering along anchored MT (red) and then MT sliding was consistently observed (Figure 2). In contrast, 3-20 nM Eg5-Δtail-GFP MT crosslinking was often observed but free-MTs (yellow) did not zipper along the anchored MT (red) and remain in scissoring motion for extensive periods (Figure 2). For MT sliding and kinesin-5 spiking studies, MT sliding experiments were performed as described above with the exception of using 20 nM FL-Eg5. Eg5-GFP spiking was carried out by the addition of either 1 nM FL-Eg5-GFP or 1 nM Eg5-Δtail-GFP with 100-200 μg/ml AlexaF-560-MTs in imaging buffer conditions. TIRF Imaging was carried out as described above.

### Image analysis of dynamic MT motility and MT sliding

Image movie stacks were preprocessed with photobleach correction and image stabilization plugins using the program FIJI (Schindelin et al., 2012). For motility along individual MTs, individual FL-Eg5-GFP or Eg5-Δtail-GFP motor motility events were identified along anchored MTs based on kymographs in generated for multiple channels. The FIJI plugin, trackmate, (Schindelin et al., 2012) was used to measure particle motility rates and identify their run lengths. Large collections of motile events for FL-Eg5-GFP or Eg5-Δtail-GFP conditions were collected for 25, 50, and 100 mM KCl conditions (Table 1). Average MT parameters were determined by frequency binning the motility events in a range conditions and then fitting these events using Gaussian distributions using the program Prism (Table 1). In general, all parameters fit single Gaussian distributions. Run lengths were fitted using exponential decay to identify the half-length for each motor condition. T-tests were performed to determine significance of the differences observed.

For motor fluorescence intensity quantifications, kymographs were manually done through analyzed using the line tool in FIJI, a line was placed over the initial signal of an individual Eg5 molecule and an intensity profile was generated and recorded in Microsoft Excel. The line was extended to include an area of the kymograph where a fluorescent signal was absent in order to measure the local surrounding background signal. This background measurement was subtracted from the initial fluorescence intensity of the molecules signal in Microsoft Excel. Only molecules that were observed to have landed on the MT during the observation period, and that were motile were used for quantification. The intensity data were frequency binned and Gaussian fit using Prism.

For MT sliding and MT sliding spiking assays, image analysis was carried out as described above with the exception of visualizing the free-MT sliding motility with respect to the anchored MT using 560 nm emission channel. The motility patterns FL-Eg5-GFP or Eg5-Δtail-GFP motor particles were studied with respect to sliding zone (along both the free and anchored MT) using the FIJI plugin, Trackmate, to determine motor motility velocities and their changes in motility direction inside or outside the sliding zone.

### Measuring pushing forces by optical trapping of Eg5 MT sliding

To study Eg5 MT sliding forces in optical trapping, flow chambers were prepared as described above and previously(Shimamoto et al., 2015). Paclitaxel stabilized Hilyte-649 and biotin labeled MTs were anchored along their surface via Biotin-Neutravidin attachment. Polystryene beads were coated with kinesin-1 nucleotide-free mutant and linked to Rhodamine labeled MTs (bead attached free-MTs). These bead attached free-MTs were then mixed with 1-20 nM FL-Eg5-GFP or 1-500 nM Eg5-Δtail-GFP and injected into these flow chambers, after being equilibrated with imaging buffer. The beads attached free MTs were observed to interact with the anchored MTs and locked into the optical trap to measure the forces. Generally, 3-10 nM FL-Eg5-GFP was sufficient to observe MT sliding events, while 3-10 nM Eg5-Δtail-GFP rarely produced sliding events, and mostly crosslinked without zippering into sliding zones forming scissoring events. At 200-500 nM Eg5-Δtail-GFP, we observed sufficient MT sliding events. For each event, the length of the sliding zone, the total Eg5-GFP intensity, and plateau pushing forces developed were measured and used for scaled comparisons (Figure 5B). The Eg5-GFP intensity scaled linearly with the MT overlap MT sliding zone length without significant difference between FL-Eg5-GFP or Eg5-Δtail-GFP conditions, despite the difference in the concentration using in the assay (Figure 5D). The plateau forces generated by FL-Eg5-GFP scaled linearly with the size of the overlap MT sliding zone length, with 9-fold difference in the slope of the same comparison for Eg5-Δtail-GFP data.

### Transfection of Hela Cells and in vivo imaging

HeLa cells were cultured in Minimal Essential Media-α (Gibco) supplemented with 10% fetal bovine serum (Gibco). Cells were maintained at 37°C with 5% CO_2_. Transient transfections of plasmid DNA were performed via electroporation using a Nucleofector 4D system, pulse code CN114, and Cell Line SE reagents (Lonza). Cells were plated onto 12 mm glass coverslips (Electron Microscopy Sciences) for fixed cell immunofluorescence, 4-chamber 35 mm glass-bottom dishes (Greiner Bio-One) for live cell imaging, or 60 mm polystyrene tissue culture dishes for lysate collection.

To assess the protein levels in vivo, cells were arrested in 100 μM monastrol (Selleckchem) overnight and lysed in PHEM buffer (60 mM PIPES, 25 mM HEPES, 10 mM EGTA, 4 mM MgSO_4_) with Halt Protease and Phosphatase Inhibitor cocktail (Thermo-Fisher) on ice. Lysates were extracted on ice for 10 minutes and centrifuged at 21,130 x g for 10 minutes. An equal volume of 4X Laemmli buffer (Bio-Rad) was added to the supernatant and samples were heated to 95°C for 10 minutes. Lysates were separated by electrophoresis on 4-15% Tris-glycine polyacrylamide gels (Bio-Rad) and transferred to polyvinylidene fluoride membranes (Bio-Rad). Membranes were blocked in Odyssey blocking reagent (LI-COR) diluted 1:1 in tris-buffered saline for 1 hour, incubated with rabbit anti-mCh (diluted 1:1000, AbCam) and mouse anti-GAPDH (diluted 1:10,000, Thermo-Fisher) primary antibodies overnight, and incubated with IRDye 800- and IRDye 680-tagged fluorescent secondary antibodies (LI-COR) for 1 hour. Blot fluorescence was imaged using an Odyssey CLx system (LI-COR) and analyzed using Image Studio Lite (LI-COR).

Fixed and live cell imaging was performed using a Nikon Ti-E inverted microscope controlled by NIS Elements software (Nikon Instruments) with a Plan APO 60X/1.42 NA oil immersion objective or APO 100X/1.49 NA oil immersion objective (Nikon Instruments), Spectra-X light engine (Lumencore), and Clara CCD camera (Andor). Image processing was performed using NIS Elements (Nikon Instruments) and ImageJ (NIH). Data analysis and statistical comparisons were performed using Excel (Microsoft) and Prism (GraphPad Software).

For assessment of mCh and Eg5-mCh expression levels and localization at metaphase in fixed HeLa cells, cells were treated with 20 μM MG132 (Selleckchem) 2 hours prior to fixation. Cells were fixed for 10 minutes in 1% paraformaldehyde (Electron Microscopy Sciences) in ice-cold methanol (Thermo-Fisher). Cells were blocked using 20% goat serum in antibody-diluting buffer (AbDil, 1X tris-buffered saline with 2% bovine serum albumin, 0.1% Triton-X 100, and 0.1% sodium azide) for 1 hour, incubated with mouse anti-α-tubulin primary antibodies (DM1a, Sigma-Aldrich, diluted 1:750 in AbDil) for 1 hour, and incubated in fluorescent secondary antibodies conjugated to Alexa Fluor 488 or 647 (Life Technologies, diluted 1:500 in AbDil) for 1 hour. Cells were mounted in ProLong Gold with DAPI (Thermo-Fisher). Expression levels of mCh-tagged proteins were compared by drawing elliptical regions of interest (ROIs) around mitotic cells using α-tubulin staining, measuring mCh fluorescence intensity within the cellular ROIs, and subtracting averaged intensity from two background ROIs containing no visible cells. Localization was assessed by defining an ROI as the spindle based on α-tubulin staining and an ROI as cytoplasm by subtracting this spindle ROI from an ellipse that encompassed the cell. Intensity of mCh signal was measured in both the spindle and cytoplasm ROIs, and a ratio of spindle to cytoplasm intensity calculated.

For live cell imaging, growth media was exchanged for CO_2_-Independent Media (Gibco) supplemented with 10% fetal bovine serum (Gibco) and penicillin/streptomycin (Gibco). For assessment of localization after treatment with BRD-9876 (Tocris Bioscience), HeLa cells were treated with 20 μM MG132 (Selleckchem) for 2 hours prior to imaging. Cells were imaged prior to drug addition, 1 minute after addition of 5 μM BRD-9876, and subsequently once every 5 minutes. For assessment of localization in anaphase, cells in metaphase were identified and imaged at 2-minute intervals through anaphase. For both live cell assays, localization of proteins to the spindle was quantified as described for fixed cell imaging, with the spindle ROI defined by GFP-tubulin signal.

## Acknowledgements

We would like to thank Dr Jonathan Scholey (Molecular Cellular Biology) for inspiring J.A.B with excitement about studying kinesin-5 mechanism. J.A.B is supported by funding support from the NSF-1615991 and NIH-GM110283. R.M. is supported by funding from NIH-GM052468. R.J.M. is supported by funding from the NIH-GM124889. S.R. is supported by funding from NIH-GM130556. L.G. is supported in part by funding from the Israel Science Foundation (ISF) (ISF 386/18), and US NSF-Israel Binational science foundation (BSF-2015851). JS is supported by funds from the NIH-GM121491 and NIH-GM130556.

## Supplementary Figure Legends

**Figure S1: The kinesin-5 tail domain inhibits the motor domain MT-activated ATPase through stabilizing the MT bound nucleotide-free state.**

A) MT co-sedimentation assays of the KLP61F motor (motor) and tail domain (tail) with MTs (tubulin) in the presence of non-hydrolysable analog, 2 mM AMPPNP at 75 mM KCl (top panel) and 25 mM KCl (bottom panel).

B) MT co-sedimentation assays of the KLP61F motor (motor) and tail domain (tail) with MTs (tubulin) in the presence of non-hydrolysable analog, 2 mM ADP.AlF4, mimicking the ADP.Pi state at 75 mM KCl (top panel) and 25 mM KCl (bottom panel).

C) MT co-sedimentation assays of the KLP61F motor (motor) and tail domain (tail) with MTs (tubulin) in the presence of the 2 mM ADP at 75 mM KCl (top panel) and 25 mM KCl (bottom panel).

D) MT co-sedimentation assays of the KLP61F motor (motor) and tail domain (tail) with MTs (tubulin) in the nucleotide-free state, achieved by adding 10 U of Apyrase. at 75 mM KCl (top panel) and 25 mM KCl (bottom panel). E) MT co-sedimentation assays of the KLP61F tail domain (tail) with MTs (tubulin) in the presence of 2 mM ADP at 75 mM KCl (top panel) and 25 mM KCl (lower panel).

F) MT co-sedimentation assays of the KLP61F motor domain (motor) with MTs (tubulin) in the presence of ADP at 75 mM KCl (top panel) and 25 mM KCl (lower panel)

G) Steady state ATP hydrolysis for Eg5 motor (blue) and equimolar Eg5-motor + tail constructs (green) with increasing MT concentrations.

H) MT co-sedimentation assays of the human Eg5 motor domain (Eg5 motor) and tail domain (Eg5 tail) with MTs (tub) in the presence of 2 mM ADP. Note the Eg5 tail domain shows a minor degradation form which also behaves in a similar manner to the full length (marked by *tail).

I) MT co-sedimentation assays of the human Eg5 motor domain (Eg5 motor) and tail domain (Eg5 tail) with MTs (tub) in the presence of 2 mM ATPgammaS.

J) Left panel quantitative densitometry of the molar ratios of Eg5 motor-tubulin (blue) and tail-tubulin (green) and motor-tubulin (left panel). These data show that the ratios are similar to the KLP61F experiments with the tail and motor to tubulin ratios being higher in the nucleotide-free and ADP states.

**Figure S2: The kinesin-5 tail domain engages the motor domain through direct interface near α0 helical hairpin in the nucleotide-free state.**

A) Top panel (I): Colores view of the KLP61F motor MT AMPPNP map, colored based on resolution scale.

Bottom Panel (II): A gold standard Fourier Shell correlation (FSC) curve for the MT decorated KLP61F motor -AMPPNP state revealing an overall resolution of 4.4 Å.

B) Top panel (I): Colores view of the KLP61F motor + tail nucleotide-free MT decorated map colored based on resolution scale. Note the small size of the tail region.

Bottom Panel (II); A gold standard Fourier Shell correlation (FSC) curve for the MT decorated KLP61F motor nucleotide free map state revealing an overall resolution of 4.4 Å.

C) Top panel (I): Colores view of the KLP61F motor + tail nucleotide-free MT decorated map after MT-patch refinement (see materials and methods). Note the increased size of the tail region but its low 8-Å resolution. The motor density resolution improved through the refinement procedure.

Bottom Panel (II); A gold standard Fourier Shell correlation (FSC) curve for the MT decorated KLP61F motor and tail in the nucleotide-free state revealing an overall resolution of 4.0 Å.

D) Class averages of the KLP61F motor AMPPNP decorated MTs (AMPPNP top), and the KLp61F motor + tail nucleotide-free state decorated MTs (Nucleotide-free bottom). Right panels, Top, a single AMPPNP motor decorated MT, compared to nucleotide free motor + tail decorated MT density is extracted and magnified. These reveal the general conformational change of the motor domain and average density for the tail around the single binding site on the backside of the motor domain.

E) An overlay of the kinesin-5 motor, three subdomain highlighted, nucleotide-free state model to the kinesin-1 ATP-like state in faded highlighted subdomains revealing the conformational change in α0 helix in kinesin-5 compared to kinesin-1 ATP state. Loops L6 and L8 are seen at unique conformation compared to kinesin-1 and are labeled with black arrows.

F) An overlay of the kinesin-5 subdomain nucleotide-free model to the kinesin-1 nucleotide-free state model revealing the nearly identical conformation of α0 helix. Loops L6 and L8 are seen at unique conformation compared to kinesin-1 and are labeled with black arrows.

G) View of the raw kinesin-5 motor tail nucleotide free map after patch refinement (as seen in panel C) with the segmentation for the map shown in the middle motor domain. Motor domain is shown in yellow, tail domain is shown in blue while α and β-tubulin are shown in cyan and green, respectively.

**Figure S3: The tail domain down-regulates the motility velocity of homotetrameric kinesin-5 motors along MTs.**

A) Right panel, Size exclusion chromatography (SEC) for recombinant FL-Eg5-GFP (red) and SDS-PAGE lane for peak fraction. Left panel, SEC for recombinant Eg5-Δtail-GFP and SDS-PAGE lane for peak fraction.

B) Two color TIRF Fields for surface anchored MTs (red) with FL-Eg5-GFP (left) and Eg5-Δtail-GFP (right) revealing the highly robust motility activity in these imaging conditions at pH 7.5 at 25-100 mM KCl.

C) Frequency distribution for motile Fl-Eg5-GFP (red) and Eg5-Δtail-GFP (blue) motor run length at 50 and 100 mM KCl. These were fit with logarithmic decay curves to determine the average run length values. The run length analyses reveal that Fl-Eg5-GFP retains processive motility in both 50 and 100 mM KCl conditions, in contrast to Eg5-Δtail-GFP which is processive at 50 mM KCl but shows a two-fold decrease in run lengths at 100 mM KCl.

D) Additional example kymographs for Fl-Eg5-GFP as shown in Figure 3D.

E) Additional example kymographs for Eg5-Δtail-GFP as shown in Figure 3E.

**Figure S4: The kinesin-5 tail domain regulates the zippering of two sliding MTs via slow directional motility within the overlapping zones.**

A) Left side panels, Wide field view of reconstituting 10-20 nM FL-Eg5-GFP mediated MT sliding events. Free MTs (yellow) can be seen recruited along anchored MTs (red) mediated by the FL-Eg5-GFP motors. Right panels, wide field image of reconstitutions of 10-20 nM Eg5-Δtail-GFP revealing a defect in MT sliding zone leading the free MT (yellow) to rotate around a single point, or scissor with respect to the anchored MT.

B) Left, Time-lapse montage/kymograph reveal how Eg5 FL-GFP motors mediates crosslinking and then zippering of newly captured MT. Left panel montage in three colors showing the capture of the free MT (yellow) by Fl-Eg5-GFP motors (green) along the anchored MT, second panel, kymograph for event in three channels. Third panel, free MT channel. Fourth panel, FL-Eg5-GFP channels. The latter three panels show the boundaries of the free-MT sliding marked with broken lines.

Right panels, Time-lapse montage/kymograph reveal how Eg5-Δtail-GFP motors mediates crosslinking and poor zippering leading to scissoring ofthe newly captured MT. Left panel, two color kymograph showing anchored MT (red) by Eg5-Δtail-GFP motors (green) along the anchored MT.

C) Additional Kymographs, similar to Figure 4F, for motor spiking into MT sliding assays. Left panels, 1 nM FL-Eg5-GFP motor (green) is spiked into MT sliding events formed by FL-Eg5 where free MT (yellow) is being slide along anchored MT (red). Note the unidirectional motility of the FL-EG5-GFP motors and their slow motility within the MT sliding zone. Right panels, 1 nM Eg5-Δtail-GFP are spiked into MT sliding events formed by FL-Eg5-GFP. Note the bi-directional motility of Eg5-Δtail-GFP within MT sliding zones.

## Supplementary Video Legends

**Video S1:** Structural transition of the Kinesin-5 motor domain from AMPPNP to nucleotide state and its effect on binding the tail domain. View of the kinesin-5 motor domain map with AMPPNP showing the motor domain model, transition to the motor nucleotide state map showing the site of binding for the tail domain density and model for the motor. Views of the two states using three motor subdomains and conformational change in the N-terminal subdomain (blue) and its effect on the ATP binding site and its rotation around the Upper subdomain (pink) and the MT bound subdomain (red).

**Video S2:** Wide view of FL-Eg5-GFP and Eg5-Δtail-GFP (green) along single MTs (Red)

**Video S3:** close up views of FL-Eg5-GFP (left) and Eg5-Δtail-GFP (right) along single MTs at 25, 50 and 100 mM KCl conditions

**Video S4:** left, close up views of FL-Eg5-GFP motors (green) mediating zippering of free MT (yellow) along anchored MT (red). Right, close up view of Eg5-Δtail-GFP motors crosslinking but unable to zipper MTs leading to scissoring defect.

**Video S5:** Top left, close up view of 1 nM FL-Eg5-GFP motor (green) spiking during free MT (yellow) sliding along anchored MT (red) mediated by 20 nM unlabeled FL-Eg5. Top right, same event without the free MT revealing FL-Eg5-GFP motors along anchored MT. bottom left, close up view of 1 nM Eg5-Δtail-GFP motor (green) spiking during free MT (yellow) sliding along anchored MT (red) mediated by 20 nM unlabeled FL-Eg5. Bottom right, same event without the free MT revealing Eg5-Δtail-GFP motors along anchored MT.

**Video S6:** left, optical trapping of MT sliding experiments revealing the bead attached to sliding free MT (red) along anchored MT (purple) mediated by FL-Eg5-GFP motors. Right, optical trapping of MT sliding experiments revealing the bead attached to sliding free MT (red) along anchored MT (purple) mediated by Eg5-Δtail-GFP motors.

## References

Acar, S., Carlson, D.B., Budamagunta, M.S., Yarov-Yarovoy, V., Correia, J.J., Ninonuevo, M.R., Jia, W., Tao, L., Leary, J.A., Voss, J.C., et al. (2013). The bipolar assembly domain of the mitotic motor kinesin-5. Nat Commun 4, 1343.

Al-Bassam, J. (2014). Reconstituting dynamic microtubule polymerization regulation by TOG domain proteins. Methods Enzymol 540, 131–148.

Blangy, A., Lane, H.A., d’Herin, P., Harper, M., Kress, M., and Nigg, E.A. (1995). Phosphorylation by p34cdc2 regulates spindle association of human Eg5, a kinesin-related motor essential for bipolar spindle formation in vivo. Cell 83, 1159–1169.

Britto, M., Goulet, A., Rizvi, S., von Loeffelholz, O., Moores, C.A., and Cross, R.A. (2016). Schizosaccharomyces pombe kinesin-5 switches direction using a steric blocking mechanism. Proc Natl Acad Sci U S A 113, E7483–E7489.

Brust-Mascher, I., Sommi, P., Cheerambathur, D.K., and Scholey, J.M. (2009). Kinesin-5-dependent poleward flux and spindle length control in Drosophila embryo mitosis. Mol Biol Cell 20, 1749–1762.

Cao, L., Wang, W., Jiang, Q., Wang, C., Knossow, M., and Gigant, B. (2014). The structure of apo-kinesin bound to tubulin links the nucleotide cycle to movement. Nat Commun 5, 5364.

Duselder, A., Fridman, V., Thiede, C., Wiesbaum, A., Goldstein, A., Klopfenstein, D.R., Zaitseva, O., Janson, M.E., Gheber, L., and Schmidt, C.F. (2015). Deletion of the Tail Domain of the Kinesin-5 Cin8 Affects Its Directionality. J Biol Chem 290, 16841–16850.

Edamatsu, M. (2014). Bidirectional motility of the fission yeast kinesin-5, Cut7. Biochem Biophys Res Commun 446, 231–234.

Forth, S., and Kapoor, T.M. (2017). The mechanics of microtubule networks in cell division. J Cell Biol 216, 1525–1531.

Fridman, V., Gerson-Gurwitz, A., Shapira, O., Movshovich, N., Lakamper, S., Schmidt, C.F., and Gheber, L. (2013). Kinesin-5 Kip1 is a bi-directional motor that stabilizes microtubules and tracks their plus-ends in vivo. J Cell Sci 126, 4147–4159.

Gerson-Gurwitz, A., Thiede, C., Movshovich, N., Fridman, V., Podolskaya, M., Danieli, T., Lakamper, S., Klopfenstein, D.R., Schmidt, C.F., and Gheber, L. (2011). Directionality of individual kinesin-5 Cin8 motors is modulated by loop 8, ionic strength and microtubule geometry. EMBO J 30, 4942–4954.

Goshima, G., and Scholey, J.M. (2010). Control of mitotic spindle length. Annu Rev Cell Dev Biol 26, 21–57.

Goshima, G., Wollman, R., Stuurman, N., Scholey, J.M., and Vale, R.D. (2005). Length control of the metaphase spindle. Curr Biol 15, 1979–1988.

He, S., and Scheres, S.H.W. (2017). Helical reconstruction in RELION. J Struct Biol 198, 163–176.

Henty-Ridilla, J.L., Rankova, A., Eskin, J.A., Kenny, K., and Goode, B.L. (2016). Accelerated actin filament polymerization from microtubule plus ends. Science 352, 1004–1009.

Heymann, J.B., and Belnap, D.M. (2007). Bsoft: image processing and molecular modeling for electron microscopy. J Struct Biol 157, 3–18.

Hildebrandt, E.R., Gheber, L., Kingsbury, T., and Hoyt, M.A. (2006). Homotetrameric form of Cin8p, a Saccharomyces cerevisiae kinesin-5 motor, is essential for its in vivo function. J Biol Chem 281, 26004–26013.

Huehn, A., Cao, W., Elam, W.A., Liu, X., De La Cruz, E.M., and Sindelar, C.V. (2018). The actin filament twist changes abruptly at boundaries between bare and cofilin-decorated segments. J Biol Chem 293, 5377–5383.

Kapitein, L.C., Kwok, B.H., Weinger, J.S., Schmidt, C.F., Kapoor, T.M., and Peterman, E.J. (2008). Microtubule cross-linking triggers the directional motility of kinesin-5. J Cell Biol 182, 421–428.

Kapitein, L.C., Peterman, E.J., Kwok, B.H., Kim, J.H., Kapoor, T.M., and Schmidt, C.F. (2005). The bipolar mitotic kinesin Eg5 moves on both microtubules that it crosslinks. Nature 435, 114–118.

Kashina, A.S., Scholey, J.M., Leszyk, J.D., and Saxton, W.M. (1996). An essential bipolar mitotic motor. Nature 384, 225.

Kwok, B.H., Kapitein, L.C., Kim, J.H., Peterman, E.J., Schmidt, C.F., and Kapoor, T.M. (2006). Allosteric inhibition of kinesin-5 modulates its processive directional motility. Nat Chem Biol 2, 480–485.

Mayer, T.U., Kapoor, T.M., Haggarty, S.J., King, R.W., Schreiber, S.L., and Mitchison, T.J. (1999). Small molecule inhibitor of mitotic spindle bipolarity identified in a phenotype-based screen. Science 286, 971–974.

Owens, B. (2013). Kinesin inhibitor marches toward first-in-class pivotal trial. Nat Med 19, 1550.

Roostalu, J., Hentrich, C., Bieling, P., Telley, I.A., Schiebel, E., and Surrey, T. (2011). Directional switching of the kinesin Cin8 through motor coupling. Science 332, 94–99.

Schindelin, J., Arganda-Carreras, I., Frise, E., Kaynig, V., Longair, M., Pietzsch, T., Preibisch, S., Rueden, C., Saalfeld, S., Schmid, B., et al. (2012). Fiji: an open-source platform for biological-image analysis. Nat Methods 9, 676–682.

Scholey, J.E., Nithianantham, S., Scholey, J.M., and Al-Bassam, J. (2014). Structural basis for the assembly of the mitotic motor Kinesin-5 into bipolar tetramers. Elife 3, e02217.

Shang, Z., Zhou, K., Xu, C., Csencsits, R., Cochran, J.C., and Sindelar, C.V. (2014). High-resolution structures of kinesin on microtubules provide a basis for nucleotide-gated force-generation. Elife 3, e04686.

Shapira, O., Goldstein, A., Al-Bassam, J., and Gheber, L. (2017). A potential physiological role for bi-directional motility and motor clustering of mitotic kinesin-5 Cin8 in yeast mitosis. J Cell Sci 130, 725–734.

Sharp, D.J., McDonald, K.L., Brown, H.M., Matthies, H.J., Walczak, C., Vale, R.D., Mitchison, T.J., and Scholey, J.M. (1999). The bipolar kinesin, KLP61F, cross-links microtubules within interpolar microtubule bundles of Drosophila embryonic mitotic spindles. J Cell Biol 144, 125–138.

Shimamoto, Y., Forth, S., and Kapoor, T.M. (2015). Measuring Pushing and Braking Forces Generated by Ensembles of Kinesin-5 Crosslinking Two Microtubules. Dev Cell 34, 669–681.

Singh, S.K., Pandey, H., Al-Bassam, J., and Gheber, L. (2018). Bidirectional motility of kinesin-5 motor proteins: structural determinants, cumulative functions and physiological roles. Cell Mol Life Sci 75, 1757–1771.

Sturgill, E.G., Norris, S.R., Guo, Y., and Ohi, R. (2016). Kinesin-5 inhibitor resistance is driven by kinesin-12. J Cell Biol 213, 213–227.

Subramanian, R., and Kapoor, T.M. (2012). Building complexity: insights into self-organized assembly of microtubule-based architectures. Dev Cell 23, 874–885.

Telley, I.A., Bieling, P., and Surrey, T. (2011). Reconstitution and quantification of dynamic microtubule end tracking in vitro using TIRF microscopy. Methods Mol Biol 777, 127–145.

Turner, J., Anderson, R., Guo, J., Beraud, C., Fletterick, R., and Sakowicz, R. (2001). Crystal structure of the mitotic spindle kinesin Eg5 reveals a novel conformation of the neck-linker. J Biol Chem 276, 25496–25502.

Vale, R.D. (2003). The molecular motor toolbox for intracellular transport. Cell 112, 467–480.

Vale, R.D., and Milligan, R.A. (2000). The way things move: looking under the hood of molecular motor proteins. Science 288, 88–95.

Valentine, M.T., Fordyce, P.M., and Block, S.M. (2006a). Eg5 steps it up! Cell Div 1, 31.

Valentine, M.T., Fordyce, P.M., Krzysiak, T.C., Gilbert, S.P., and Block, S.M. (2006b). Individual dimers of the mitotic kinesin motor Eg5 step processively and support substantial loads in vitro. Nat Cell Biol 8, 470–476.

van den Wildenberg, S.M., Tao, L., Kapitein, L.C., Schmidt, C.F., Scholey, J.M., and Peterman, E.J. (2008). The homotetrameric kinesin-5 KLP61F preferentially crosslinks microtubules into antiparallel orientations. Curr Biol 18, 1860–1864.

von Loeffelholz, O., and Ann Moores, C. (2019). Cryo-EM structure of the Ustilago maydis kinesin-5 motor domain bound to microtubules. J Struct Biol.

von Loeffelholz, O., Pena, A., Drummond, D.R., Cross, R., and Moores, C.A. (2019). Cryo-EM Structure (4.5-A) of Yeast Kinesin-5-Microtubule Complex Reveals a Distinct Binding Footprint and Mechanism of Drug Resistance. J Mol Biol.

Wang, H., Brust-Mascher, I., and Scholey, J.M. (2014). Sliding filaments and mitotic spindle organization. Nat Cell Biol 16, 737–739.

Weinger, J.S., Qiu, M., Yang, G., and Kapoor, T.M. (2011). A nonmotor microtubule binding site in kinesin-5 is required for filament crosslinking and sliding. Curr Biol 21, 154–160.

Zhang, K. (2016). Gctf: Real-time CTF determination and correction. J Struct Biol 193, 1–12.

