## Supplementary material for "The Kinesin-5 Tail Domain Directly Modulates the Mechanochemical Cycle of the Motor for Anti-Parallel Microtubule Sliding": tables S1-S4

**Table 1**: steady kinetic parameters for MT activated ATP hydrolysis

| **Construct** | **Source** | **Salt con** | **k_cat_ (sec^-1^)** | **K_0.5,MT_ (nM)** |
| --- | --- | --- | --- | --- |
| Motor | *Dm KLP61F* | 50 mM KAc | 7.1 ± 0.1 | 680 ± 48 |
| Motor + Tail | *Dm KLP61F* | 50 mM KAc | 3.5 ± 0.5 | 757 ± 327 |
| Motor-Tail fusion | *Dm KLP61F* | 50 mM KAc | 3.3 ± 0.1 | 39 ± 11 |
| Motor | Hs Eg5 | 20 mM KCl | 7.3 ± 0.2 | 84 ± 14 |
| Motor + Tail | Hs Eg5 | 20 mM KCl | 5.4 ± 0.3 | 209 ± 56 |

**Table 2**: Cryo-EM KLp61F motor and tail MT structures: collection and reconstruction

|  | Dm KLP61F motor-AMPPNP  (15 protofilaments) | Dm KLP61F motor AMPPNP  (14 protofilaments) | Dm KLP61F motor+ tail- rigor  (15 protofilaments) | Dm KLP61F5 motor-tail-rigor  (14-protofilaments) |
| --- | --- | --- | --- | --- |
| **Data collection** |  |  |  |  |
| Microscope | Titan Krios (FEI) | Titan Krios (FEI) | Titan Krios (FEI) | Titan Krios (FEI) |
| Voltage (kV) | 300 | 300 | 300 | 300 |
| Ls | 22,500X | 22,500X | 22,500X | 22,500X |
| Cumulative exposure dose (e^-^ Å^-2^) | 38 | 38 | 40 | 40 |
| Exposure rate (e^-^/pixel/sec) | 7.9 | 7.9 | 8.3 | 8.3 |
| Detector | K2 Summit | K2 Summit | K2 Summit | K2 Summit |
| Pixel size (Å)* | 1.31 | 1.31 | 1.31 | 1.31 |
| Defocus range (µm) | 0.3-3.78 | 0.7-3.78 | 0.19-5.12 | 0.19-5.12 |
| Average defocus (µm) | 1.75 | 1.75 | 1.86 | 1.86 |
| Micrographs Used | 1260 | 1260 | 955 | 955 |
| Total extracted helical segment (no.) | 73,451 | 73,451 | 44,081 | 44,081 |
| Refined helical segment (no.) | 21,004 | 39,001 | 9,490 | 27,433 |
| **Reconstruction** |  |  |  |  |
| Final helical segments (no.) | 21,004 | 39,001 | 9,490 | 27,433 |
| Symmetry imposed | HP | HP | HP | HP |
| Resolution (global) FSC 0.143 | 4.2 | 4.4 | 4.2 | 4.5 |

**Table 3: Motility parameters for FL-Eg5-GFP and Eg5-Δtail-GFP along single MTs**

| **FL-Eg5-GFP** | **velocity (nm/s)** | **Motor Fluorescence (Au)** | **Run length (μm)** |
| --- | --- | --- | --- |
| 25 mM KCl | 7 ± 0.5 n=149 | 1080± 30 n=100 | N/A |
| 50 mM KCl | 26 ± 4 n=200 | N/A | 13.7 ±0.6 |
| 100 mM KCl | 26 ± 5 (60%)  41 ± 4 (40%) n=149 | 2277 ± 100  4467 ± 630 n=92 | 13.08±0.6 |
| **Eg5-Δtail-GFP** |  |  |  |
| 25 mM KCl | 32 ± 5 n=421 | 960 ± 20 n=95 | N/A |
| 50 mM KCl | 33 ± 4 n=420 | N/A | 14.9±0.6 |
| 100 mM KCl | 36 ± 5 (85%)  55 ± 10 (15%) n=149 | 1450 ± 3 n=95 | 8.0 ± 0.6 |

**Table 4: Motor motility and MT sliding motility parameters *in vitro* MT sliding assays**

| Single motor velocities in relation to free MT sliding motility | | |
| --- | --- | --- |
| **FL-Eg5-GFP** | **Free MT sliding motility (nm/s)** | **Motility in sliding zones (nm/s)** |
| **25 mM KCl** | 13.8 ± 1.0 n=26 | 13.9 ± 1.0 n=32 |
| **50 mM KCl** | 31.2 ± 1.2 n=33 | 22.7 ± 1.2 n=71 |
| Single motor motility within MT sliding zones | | |
|  | **Eg5-Δtail-GFP motors (nm/s)** | **FL-Eg5-GFP motors (nm/s)** |
| **Inside zone** | 8.6 ± 0.9 n=32 | 3.4 ± 0.3 n=67 |
| **Outside zone** | 9.6 ± 0.8 n=52 | 5.6 ± 0.3 n= 45 |
