## Supplementary figures and images for "The Kinesin-5 Tail Domain Directly Modulates the Mechanochemical Cycle of the Motor for Anti-Parallel Microtubule Sliding"

### Figure S1

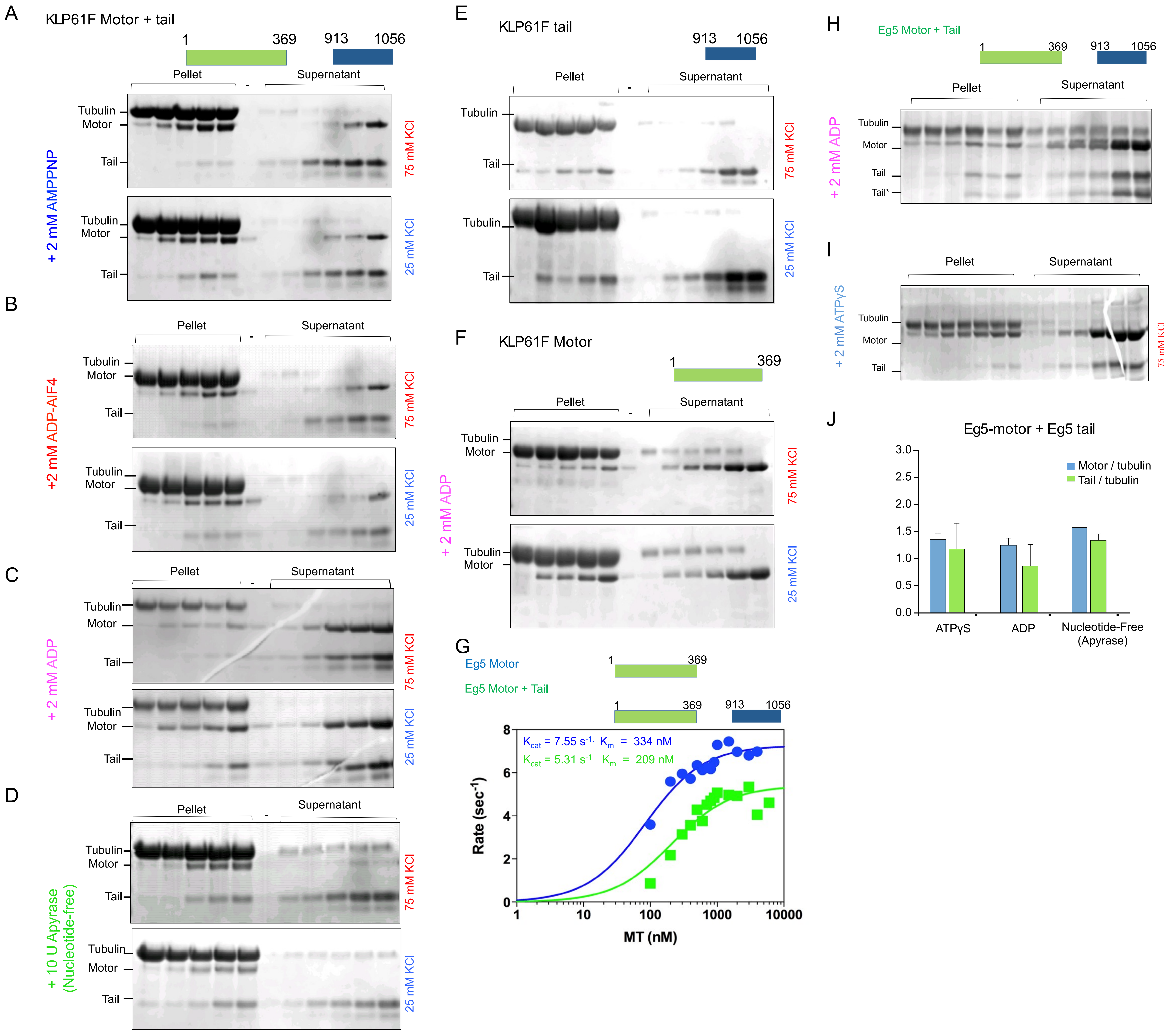

### Figure S2

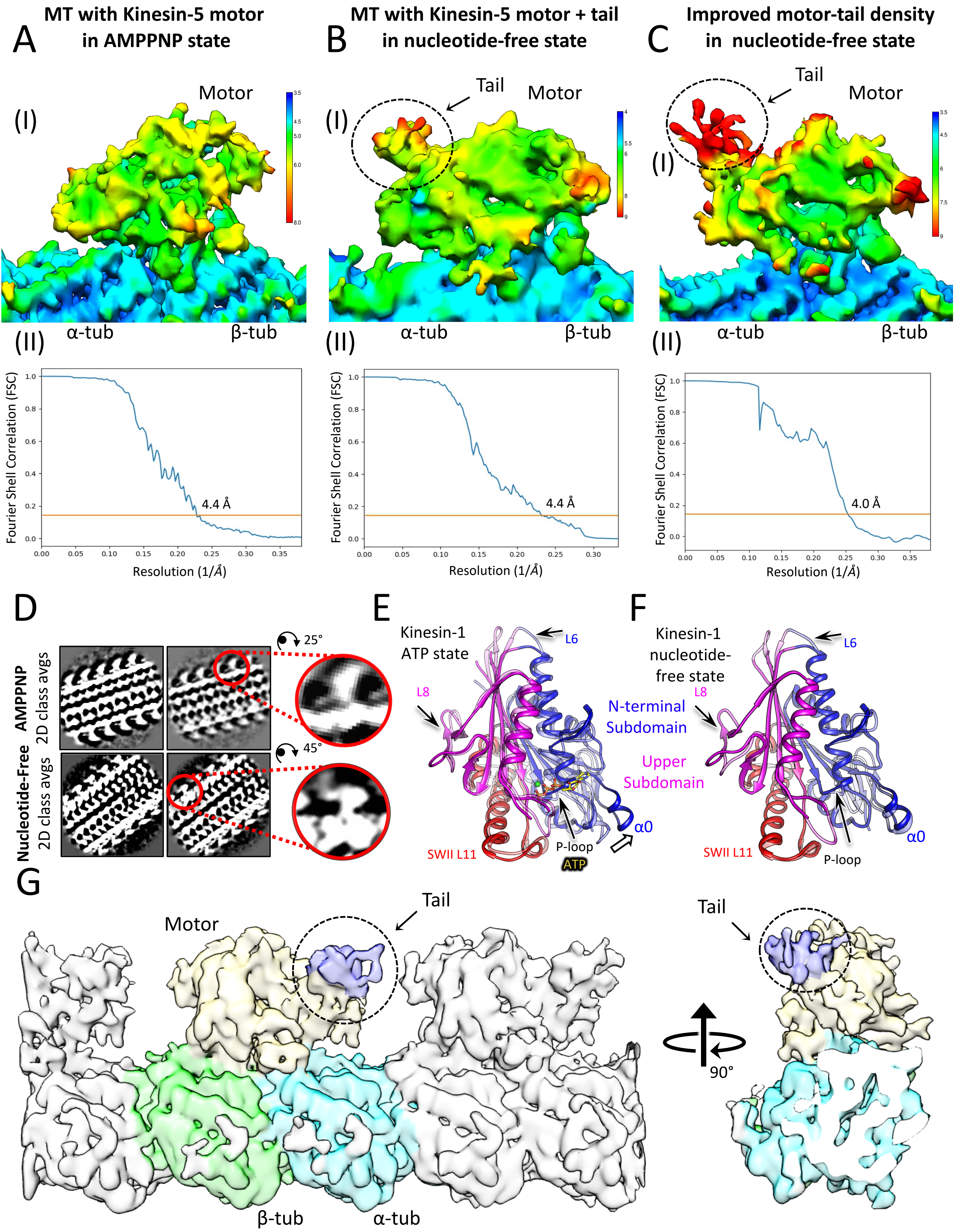

### Figure S3

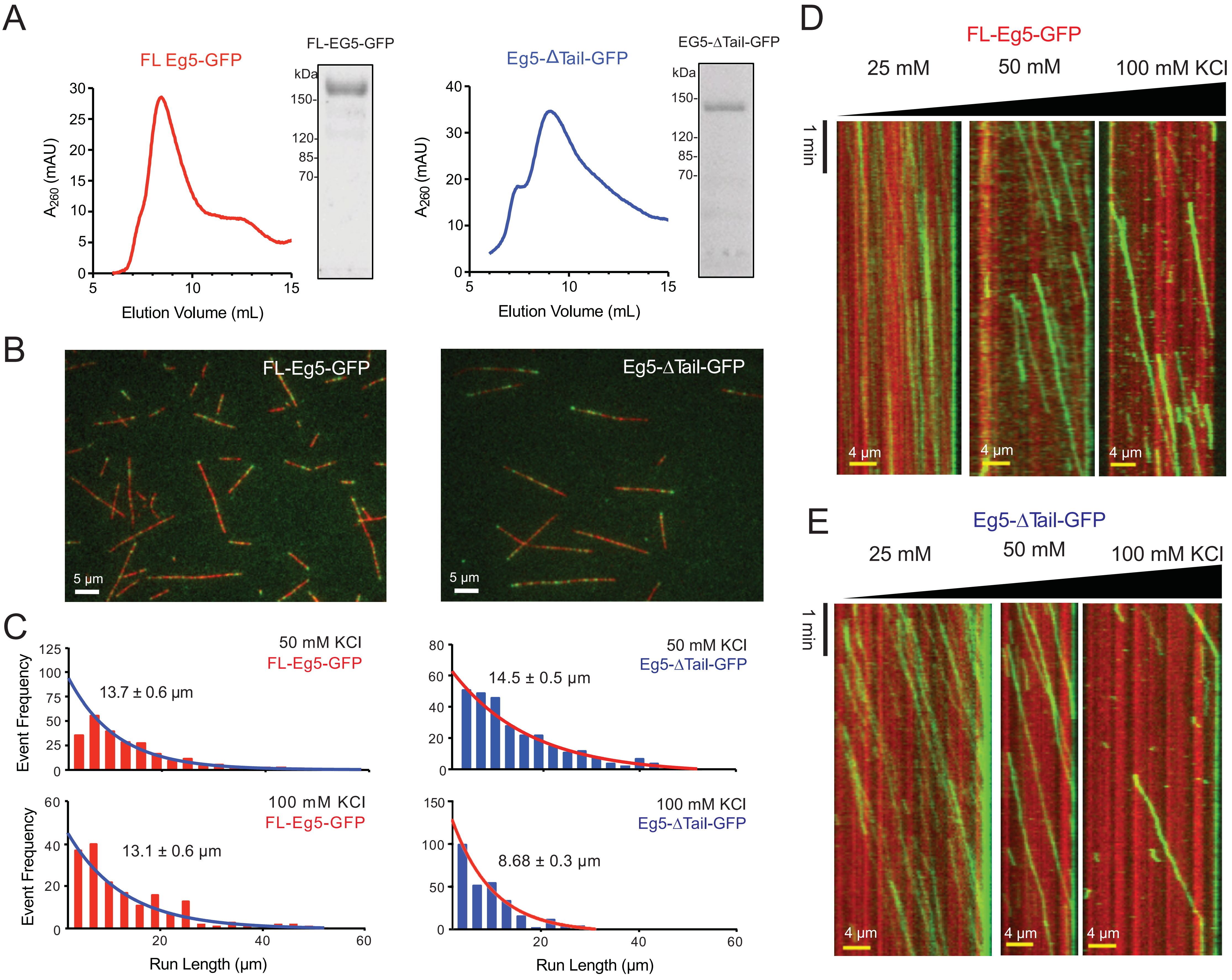

### Figure S4

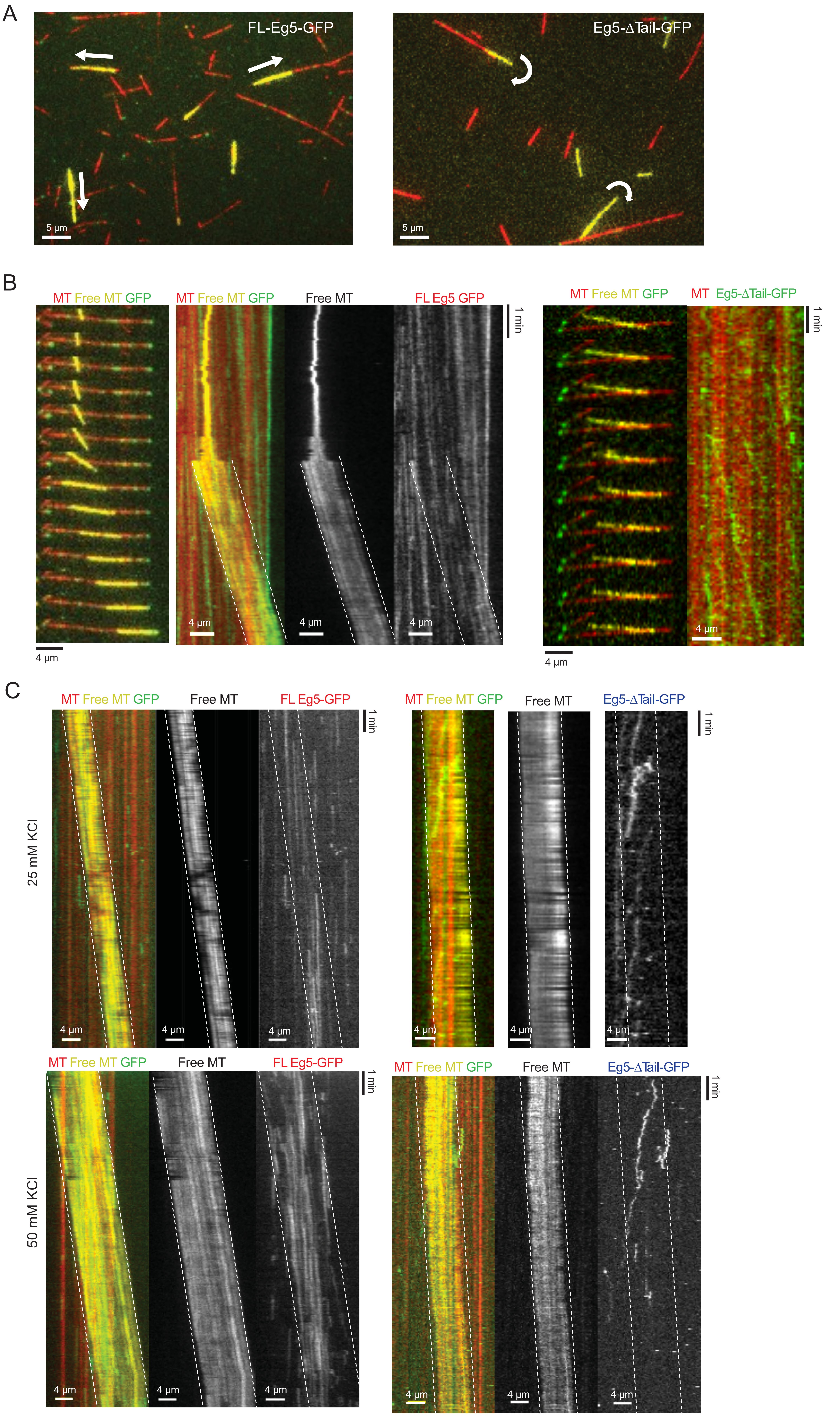
